# A single Omicron mutation reshapes ORF3a-driven host-cell remodelling

**DOI:** 10.64898/2026.08.02.742305

**Authors:** Ana de Lucas-Rius, Miguel Padilla-Blanco, Unai Merino-Herrán, Laura Mendoza-García, Ana J. Pérez-Berná, Blanca D. López-Ayllón, Camila Scagnetti Zambrano, Juozas Grigas, Oihane E. Albóniga, Jorge Montesinos, Francisco J. Chichón, Oscar Fernández, Kevin Mamprin, Raúl Fernández-Rodríguez, Juan M. Falcón-Pérez, Noa B. Martín-Cófreces, Tránsito García-García, Juan J. Garrido, María A. Oliva, María Montoya

## Abstract

SARS-CoV-2 ORF3a remodels host membranes, but the structural basis and metabolic consequences of this process remain unclear. Here, we combine complementary imaging approaches to define ORF3a function at nanometric scale, identifying underlying mechanisms, and determining how Omicron variant rewire this activity. ORF3a from the ancestral Wuhan strain disrupts Golgi cisternae, drives the formation of ORF3a dense vesicles, remodels mitochondrial architecture, and promotes lipid droplet expansion. Multi-omics analyses further reveal selective triacylglycerol accumulation linked to DGAT1 upregulation, which we validate pharmacologically through DGAT1 inhibition. In contrast, Omicron ORF3a variant, despite carrying only the Thr223Ile substitution within the β7-β8 loop at the bottom of the ‘cytosolic domain’, induced a dramatic phenotypic shift: ORF3a localizes to multivesicular bodies, preserves Golgi architecture, and fails to induce lipid accumulation. All together, these results identify ORF3a as a regulator of membrane organization and lipid homeostasis, showing how minimal sequence variation rewires host-cell remodelling.

**Graphical TOC:** 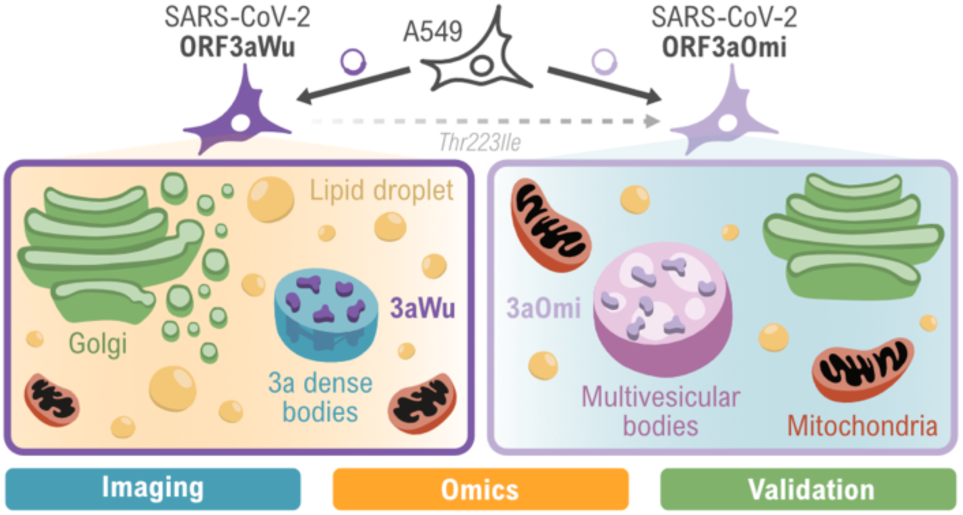

## INTRODUCTION

Since its emergence in late 2019, SARS-CoV-2 has driven intense efforts to determine the molecular determinants underlying COVID-19 pathogenesis. Initial studies focused on the four structural proteins – spike (S), envelope (E), membrane (M) and nucleocapsid (N) – with particular emphasis on S due to its role in ACE2-mediated entry and its central relevance for vaccines and therapeutic interventions^1,2^. More recently, attention has shifted towards accessory proteins, including ORF3a, ORF6, ORF7a/b, ORF8 and ORF9b, which although dispensable for replication, act as critical modulators of host-pathogen interactions^3,4^.

ORF3a is the largest SARS-CoV-2 accessory protein and comprises 275-amino acids. Structurally, it forms a conserved dimer in which each protomer contains three transmembrane helices surrounding a central cavity and a ‘cytosolic’ β-sheets-rich domain capable of mediating higher-order oligomerization^5,6^. Early studies proposed that ORF3a functions as a viroporin based on its structural features and ion-flux assays. However, subsequent electrophysiological analysis challenged this interpretation and instead suggested alternative activities, including a possible role as a lysosomal water transporter^6–8^. Consistent with its broad cellular distribution, ORF3a has been implicated in multiple host processes, including inflammasome activation, interferon suppression, apoptosis, mitochondrial dysfunction, and metabolic regulation^3^. More recent work has also shown that ORF3a palmitoylation contributes to viral pathogenesis by suppressing TRIM16-dependent K27-linked ubiquitination^9^. Together, these findings indicate that the molecular functions of ORF3a remain incompletely understood and are likely to be highly context-dependent.

ORF3a is particularly recognised as a potent regulator of intracellular membrane trafficking. Studies using early SARS-CoV-2 isolates genetically related to the Wuhan strains demonstrated that ORF3a interacts with the Homotypic Fusion and Protein Sorting (HOPS) complex^10^. ORF3a disrupts autophagic flux by sequestering the HOPS subunit VPS39 through a C-terminal YXXΦ motif located in the β2-β3 loop of its ‘cytosolic domain’. This prevents RAB7A-dependent recruitment of fusion machinery and thereby blocks autophagosome-lysosome fusion^11–13^. In parallel, SARS-CoV-2 induces the formation of double-membrane vesicles (DMVs) that support viral genome replication, and ORF3a-mediated inhibition of autophagy may contribute to their persistence^12^. ORF3a also remodels membranes derived from the trans-Golgi network and endosome compartments, generating electron-dense vesicles termed “3a dense bodies” (3DBs). These structures undergo dynamic fusion and fission events and have been proposed to facilitate S-protein processing and virion maturation^14^.

Reprogramming of host lipid metabolism is a hallmark of many RNA viruses. Lipid droplets (LD) are endoplasmic reticulum (ER)-derived organelles that integrate lipid storage, cellular metabolism and immune signalling, and are frequently co-opted to support viral replication^15–18^. SARS-CoV-2 infection causes marked alterations in lipid homeostasis that correlates with disease severity, including reduced circulating lipid levels and intracellular LD accumulation in both patient-derived samples and *in vitro* systems^19–21^. ORF3a has been identified as a key driver of LD accumulation, and specific residues within its ‘cytosolic domain’ (S171 in β3, W193 in the β5-β6 loop and L219 in the β7-β8 loop) are required for this phenotype^22^. These observations suggest that the membrane-remodelling and metabolic activities of ORF3a may be mechanistically interconnected.

The emergence of SARS-CoV-2 variants further highlights the potential functional relevance of ORF3a. Although Omicron variant exhibits extensive mutations in the S protein^23^, ORF3a has remained comparatively conserved. Nevertheless, some recent Omicron lineages have acquired additional substitutions within the transmembrane region, including K67N in the KP.2.3 lineage and T89I in the MV.1 lineage. In contrast, the T223I substitution within the cytoplasmic β7-β8 loop has remained highly conserved since its emergence with the BA.2 lineage in early 2022, being retained across subsequent Omicron lineages^3,22^. Importantly, T223I has been associated with reduced LD accumulation and attenuated autophagy modulation, consistent with the lower pathogenicity associated to Omicron infections^22^. Nevertheless, structural and mechanistic effects of the T223I mutation in the Omicron variant have not yet been fully elucidated.

Here, we combine complementary imaging approaches, including transmission electron microscopy (TEM), cryo-electron tomography (cryo-ET), confocal immunofluorescence microscopy, cryo-soft X-ray tomography (cryo-SXT) and cryo-structured illumination microscopy (cryo3DSIM) in 3D, to define the multiscale structural landscape induced by ORF3a expression. These approaches reveal extensive remodelling of intracellular membranes, including the accumulation of aberrant vesicular structures, alterations in mitochondrial morphology, and pronounced LD enrichment. Integrating RNA-sequencing and lipidomic analyses further uncovers a coordinated metabolic rewiring driven by ORF3a. Mechanistically, we show that ORF3a from the Wuhan strain promotes LD biogenesis through DGAT1-dependent triacylglycerol synthesis, whereas the T223I Omicron variant markedly attenuates this activity. Together, our results established ORF3a as a major regulator of host membrane organisation and lipid homeostasis, and provide a mechanistic framework linking nanoscale membrane remodelling to metabolic reprogramming and viral pathogenicity.

## RESULTS AND DISCUSSION

### ORF3a Wuhan disrupts Golgi architecture and drives the formation of ORF3a dense vesicles

To define how ORF3a remodels host membranes, A549 cells stably expressing ORF3a from the ancestral Wuhan strain (hereafter ORF3aWu) were established **(Figure S1)**. Two types of constructs were used: a C-terminal 2×Strep-tag (ST) and a C-terminal enhanced green fluorescent protein (GFP) fusion, and included cells expressing ST or GFP alone as controls. Both types of ORF3aWu constructs produced indistinguishable phenotypes when tested and therefore, they were used interchangeably.

TEM imaging of 70 nm cell sections revealed a marked reorganization of the intracellular membrane landscape upon ORF3aWu expression. Control cells showed a typical cytoplasmic organization with dispersed mitochondria, lysosomes, and small vesicle compartment **(Figure 1A)**. In contrast, ORF3aWu cells accumulated large membranous vesicles filled with dense, disordered luminal material **(Figure 1B)** that resembled the 3DB morphology recently described^14^. This landscape is consistent with extensive endomembrane remodelling in SARS-CoV-2-infected cells, where ER-and Golgi-derived membrane proliferation leads to the accumulation of single-and double-membrane vesicles that support viral replication^24,25^. Using cryo-ET, tomograms of 250-nm-thick sections were acquired, partially resolving the three-dimensional internal organization of these compartments while preserving cellular architecture in a state as close as possible to its native condition. Thus, our result suggested that electron-dense material formed an interconnected tubular network rather than amorphous aggregates **(Figure 1C and Movie S1)**. This network resembled virus replication organelles, which have previously been reported to be enhanced by ORF3a^26^.

**Figure 1.**
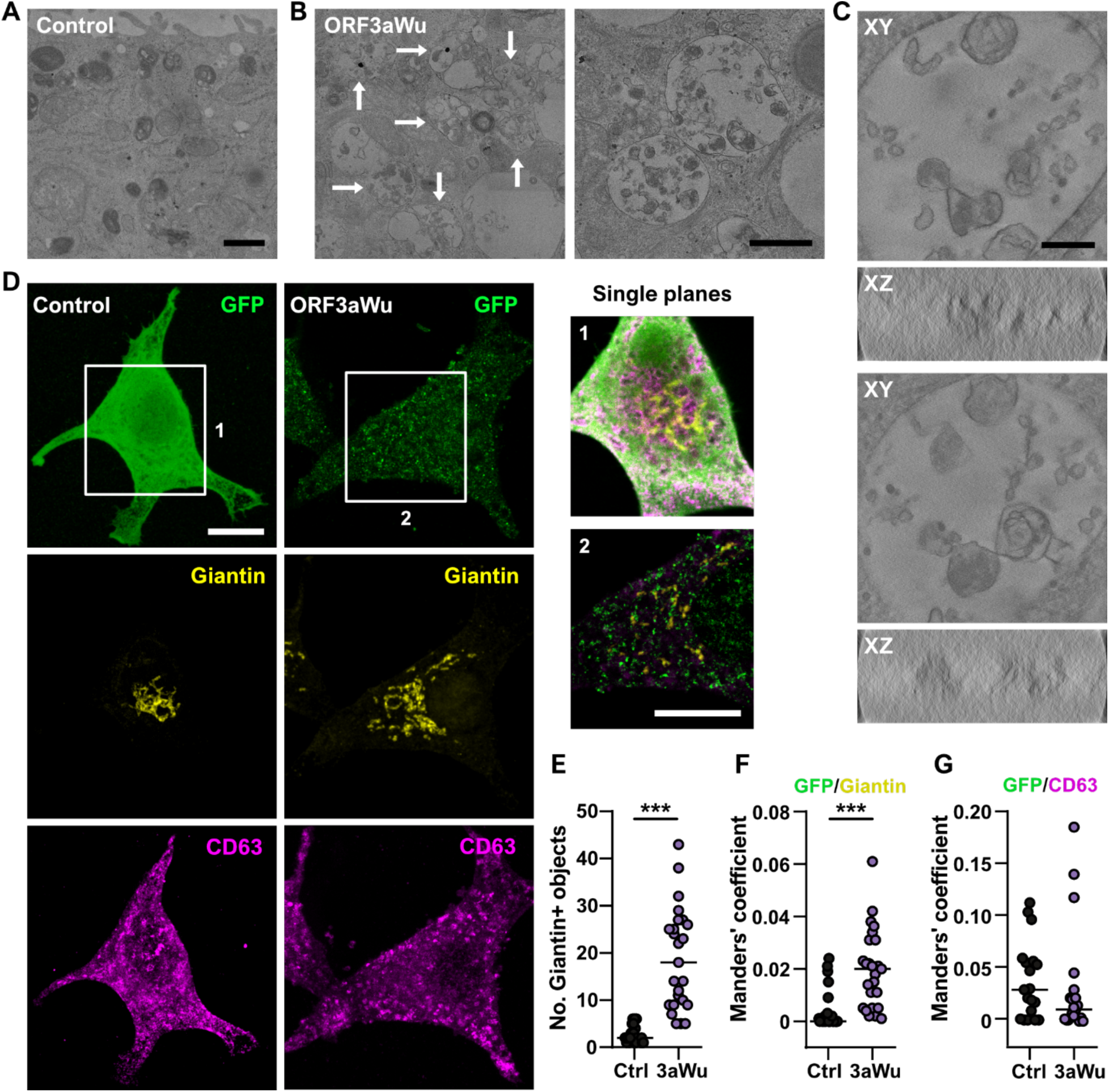
ORF3a Wuhan promotes cytosolic vesiculation and Golgi disruption. **(A-B)** TEM images of 70 nm sections of control **(A)** and ORF3aWu **(B)** A549 cells. Scale bar: 1 µm. White arrows indicate dense vesicles. On the right **(B)**, higher magnification of dense vesicles. Scale bar: 800 nm. **(C)** Two different XY and XZ planes from a 250 nm section cryo-ET of A549 ORF3aWu cell. Scale bar: 100 nm. **(D)** Maximal projections from confocal stacks showing GFP (green), Golgi (Giantin, yellow) and MVB (CD63, magenta) in control GFP or ORF3aWu-GFP A549 cells. Insets 1 and 2 are shown in right panels as overlapped and magnified, single plane images. Scale bar: 10 μm. **(E)** Quantification of Golgi particles (number of Giantin-positive objects) in control and ORF3aWu cells. **(F-G)** Mander’s coefficient for GFP co-localization with Giantin **(F)** and CD63 **(G)** in control and ORF3aWu cells. Graphs, median (control, n=28; ORF3aWu, n=25). Statistical significance is given as: ***p < 0.001

We next investigated whether ORF3aWu alters Golgi organization and whether these vesicles relate to endolysosomal compartments. Confocal imaging of cells immunostained for Giantin (Golgi) and CD63 (late endosomes and multivesicular bodies (MVB))^27^ revealed a clear disruption of Golgi architecture **(Figure 1D-G)**. Control cells showed a compact perinuclear Golgi ribbon, whereas ORF3aWu-GFP fragmented the Golgi into disperse puncta, consistent with cisternal breakdown **(Figure 1D, E)**. Previous studies reported similar Golgi disorganization in SARS-CoV-2–infected A549-ACE2 cells at late stages of infection, as well as after ORF3aWu expression alone^14,28^. Interestingly, disassembly of the trans-Golgi network as well as disrupted endosome trafficking have been described to trigger NLRP3 inflammasome activation^29,30^, including influenza viral infection^31^, and thus, ORF3a-mediated Golgi disruption may directly couple membrane remodelling to inflammatory signalling.

However, ORF3a has not been shown to be the principal accessory protein responsible for NLRP3-driven disruption when expressed alone^32^ and, during viral infection, synergistic interactions among multiple viral proteins are likely to contribute to the activation of inflammation. Quantitative analysis of ORF3aWu A549 cells showed an increase in surface area occupied by Giantin-positive structures without changes in total Golgi volume **(Figure S2)**, indicating fragmentation rather than bulk Golgi expansion. ORF3aWu puncta frequently appeared adjacent to, or partially overlapped with, Giantin-positive elements, and displayed increased colocalization compared to controls **(Figure 1F)**. Previous studies have demonstrated that ORF3a partially localizes to the Golgi, ER and endocytic compartments in infected cells^6–8,11^. This localization may be attributed to the lipid-binding sites within its transmembrane region, which likely facilitate membrane association^5,6^. Together, these results indicated that ORF3aWu alone directly perturbed Golgi architecture and promoted vesicular structures at the Golgi interface. Interestingly, despite being 73% similar to its SARS-CoV-2 homolog, SARS-CoV-1 ORF3a does not fragment Golgi network^14^, indicating that the Golgi-disrupting activity is not an intrinsic feature of ORF3a but rather a SARS-CoV-2-specific adaptation that likely arises from specific sequence or structural differences in the SARS-CoV-2 protein. At confocal resolution, ORF3aWu-GFP showed minimal overlap with CD63 **(Figure 1G)**, implying that these structures did not correspond to late endosomes and canonical MVB, supporting recent work showing that 3DBs derive from Golgi and early endosomal membranes and remain distinct from classical MVBs^14^.

To investigate the ultrastructural organization of our dense vesicles while minimizing artifacts associated with chemical fixation, staining, and resin embedding, a cryo-correlative workflow was implemented preserving cells in a near-native state. Cells were grown to ∼ 70% confluence on gold TEM grids, labelling lysosomes with LysoTracker^TM^, and samples were vitrified via rapid plunge-freezing. ORF3aWu-GFP and lysosomal signals were localized by cryo-SIM at the ALBA synchrotron, achieving super-resolution imaging of the GFP and red fluorescence channels. Subsequently, the same grids were transferred to the BL09-MISTRAL beamline for cryo-SXT, which provided detailed 3D ultrastructural information of the same regions^33^. Such correlative approach allowed us to localize ORF3aWu-GFP into complex multivesicular membrane assemblies, which may be related to previously described 3DBs **(Figure 2A, Movie S2)** and clearly different from lysosomes **(Figure 2B, Movie S3)**. A prior cryo-SXT study identified similar perinuclear vesicle clusters with electron-dense luminal content in SARS-CoV-2 infected A549-ACE2 cells at late stages of infection, further supporting this interpretation^28^. In addition, our data showed partial spatial association with lysosomes **(Figure 2C, Movie S4)**, consistent with previous reports linking ORF3a to endolysosomal trafficking pathways^6,11,12,34^. Pearson’s correlation coefficient (PCC) between ORF3aWu-GFP and lysosomes channels was 0.34, indicating a moderate positive correlation between the signal intensities. Thresholded Manders’ coefficients were tM1 = 0.339 and tM2 = 0.471, indicating that approximately 34% of the signal of lysosomes colocalized with ORF3aWu-GFP, whereas 47% of the signal of ORF3aWu-GFP colocalized with lysosomes. The Costes significance test yielded a p-value of 1, demonstrating that our observed colocalization was significantly greater than expected by chance. Overall, these results indicate a statistically significant but partial colocalization between the ORF3aWu-GFP in the lysosomes. Together, these results show that ORF3aWu disrupted Golgi cisternae and drove dense bodies vesicle formation, membrane compartments positioned at the Golgi-endolysosomal interface, which were distinct from classical CD63-positive MVB population.

**Figure 2.**
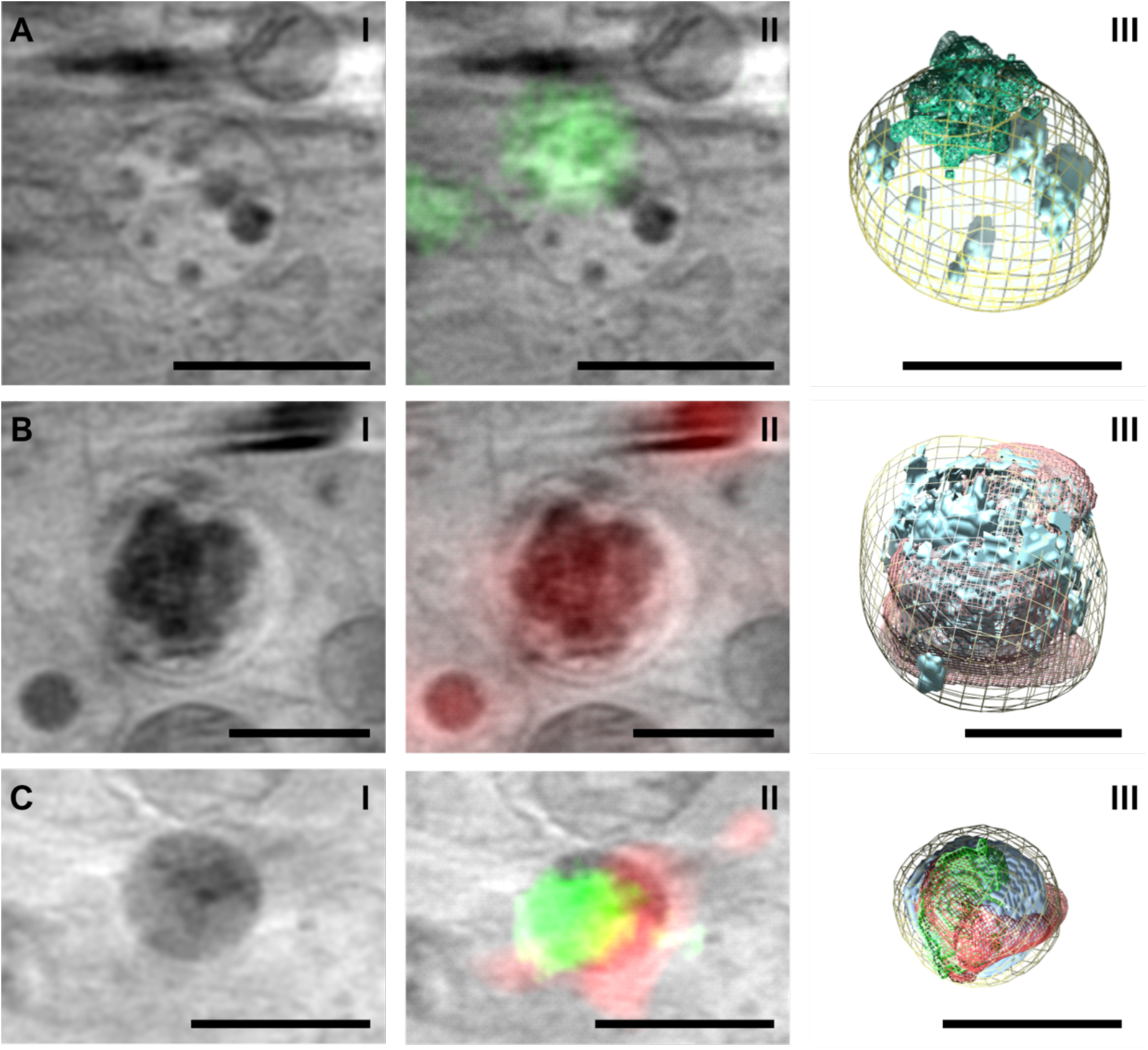
Correlative cryo3DSIM and cryo-SXT analysis of A549 cells expressing ORF3aWu-GFP. Representative examples of ORF3a dense vesicles **(A)**, lysosomes **(B)**, and vesicles positive for both ORF3aWu-GFP and LysoTracker **(C)**. For each example, the three columns show: **(I)** virtual slices through the Cryo-SX tomograms; **(II)** the corresponding tomographic slide overlaid with cryo3DSIM fluorescence, allowing the identification of ORF3aWu-GFP (green) and LysoTracker-positive structures (red); and, (**III**) manual segmentation of the vesicles (yellow) and their luminal high-contrast structures (light blue), together with the corresponding 3D ORF3aWu-GFP (green) and LysoTracker (red) fluorescence signals. Scale bars: 1 μm

### ORF3a Wuhan induces mitochondrial remodelling and lipid droplet expansion

In A549 cells, our previous results showed that ORF3aWu expression reduced mitochondria size, disrupted cristae organization, and decreased mitochondrial membrane potential (ΔΨm), consistent with metabolic dysregulations^35^. To characterize these alterations in three dimensions, cryo-epifluorescence microscopy and cryo-SXT were combined. Cryo-epifluorescence imaging identified ORF3aWu-positive regions displaying a punctuated distribution consistent with vesicular localization, in contrast to the diffuse GFP signal observed in control cells **(Figure 3A)**. These results were consistent with those shown in **Figure 1D**. Cryo-SXT reconstructions of these regions revealed a densely packed cytoplasm enriched in vesicular compartments, including electron-dense structures resembling the previously identified 3DBs^14^, which were absent from control GFP-expressing A549 cells **(Figure 2** and **3A, Movie S5 and S6,** blue organelles in segmented tomograms).

**Figure 3.**
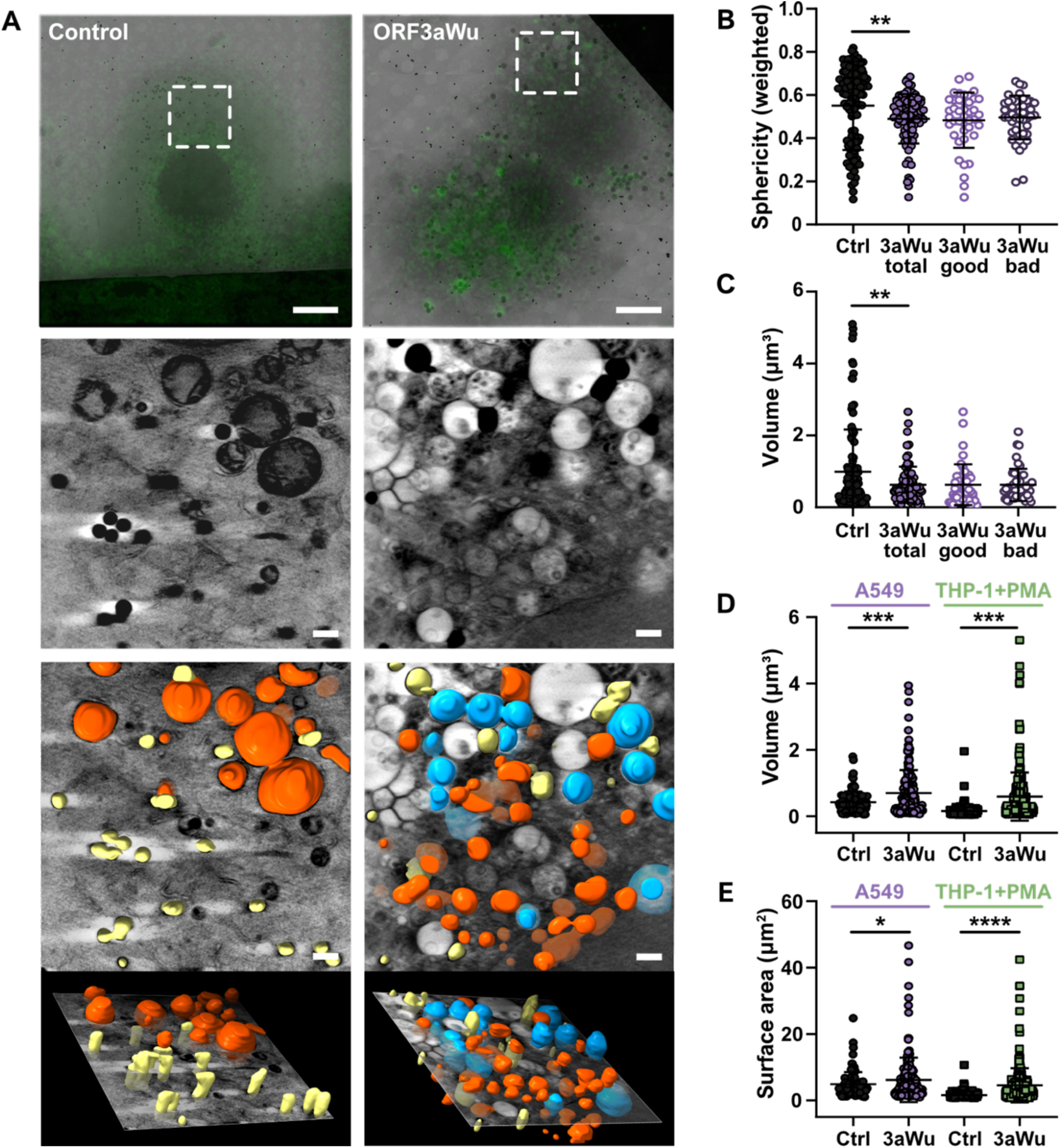
ORF3aWu transforms A549 cellular landscape. **(A)** 3D reconstruction of control and ORF3aWu-expressing A549 cells by cryo-SXT. Top panels show GFP cryo-epifluorescence signal (green, scale bar: 10 μm). Dashed boxes indicate regions of tomogram acquisition (middle panels, scale bar: 1 μm). Bottom panels show the corresponding tomograms segmentations onto representative slices of the tomograms, highlighting LD (yellow), mitochondria (red) and dense vesicles (blue) (scale bar: 1 μm). **(B)** Mitochondrial sphericity (weighted) and **(C)** volume (µm³) in control and ORF3aWu-expressing A549 cells. Total mitochondria in ORF3aWu A549 cells (“3aWu total”) were classified as normal (“3aWu good”) and altered (“3aWu bad”) according to their electron density and cristae’s appearance. **(D)** LD volume (µm³) and **(E)** surface area (µm²) in controls, ORF3aWu-expressing A549 cells and PMA-differentiated THP-1 macrophages. Data are represented as mean ± SD (n = 8 tomograms). Statistical significance is as follows: *p < 0.05, **p < 0.01, ***p < 0.001, ****p < 0.0001.

Within this extensively remodelled intracellular landscape, mitochondria **(Figure 3A**, **Movie S6,** red organelles in segmented tomograms) displayed striking structural alterations. ORF3aWu-expressing cells contained smaller mitochondria, with ∼ 50% displaying irregular morphology together with cristae abnormalities, including non-parallel and dense cristae indicative of mitochondria stress^36^ **(Figure S3A)**. Quantitative analysis of 3D reconstruction demonstrated a significant reduction in both mitochondrial volume and sphericity compared with control cells (**Figure 3B, C and S3A)**. Rather than defining distinct morphological subclasses, these alterations affected all mitochondrial network globally. Consistent with these structural alterations, MitoView staining identified a subpopulation of ORF3aWu-expressing cells with reduced ΔΨm compared to control cells **(Figure S3B, C)**. These observations closely resemble those reported in SARS-CoV-2-infected ACE2-A549 cells, in which abnormal mitochondria progressively accumulate during infection, displaying reduced size, altered cristae organization, and increased absorption contrast, ultimately representing up to 93% of the mitochondrial population at late infection stages^28^.

ORF3aWu expression also induced a marked accumulation of LDs. Cryo-SXT revealed enlarged LDs frequently positioned in close proximity to both mitochondria and vesicular compartments **(Movie S5, Figure 3A**, yellow organelles in segmented tomograms), suggesting coordinated remodelling of lipid metabolism and intracellular membrane trafficking. Quantitative analysis confirmed a significant increase in both LD volume and surface area compared with control cells **(Figure 3D, E)**. To determine whether this phenotype was restricted to lung epithelial cells, which naturally accumulate neutral lipids for surfactant production, we expressed ORF3aWu-GFP in phorbol 12-myristate 13-acetate (PMA)-differentiated THP-1 macrophages. These cells similarly accumulated enlarged and abundant LDs, closely recapitulating the phenotype observed A549 cells **(Figure 3D, E, and Figure S4)**.

LD accumulation is a hallmark of SARS-CoV-2, but not SARS-CoV-1 infection^21^, and increased LD abundance has been reported in both type II pneumocytes and circulating monocytes from patients with COVID-19^19–21^. More recently, elevated hepatic LD accumulation was observed in MA10-infected mice despite the absence of detectable viral replication in the liver, suggesting that systemic metabolic alterations may result from ORF3a activity rather than direct infection of hepatocytes. This phenotype was proposed to arise from ORF3a-mediated inhibition of autophagosome–lysosome fusion and the consequent impairment of lysosomal function, leading to defective LD turnover. Such findings demonstrate that ORF3a may induce abnormal hepatic metabolism in a mild COVID-19 model, even in the absence of detectable SARS-CoV-2 replication within the liver^37^. Consistent with previous research^22,26^, we demonstrate here that expression of ORF3aWu alone is sufficient to drive robust LD accumulation. Although ORF3a has been reported to localize around lipid LDs based on immunofluorescence analyses^26^, our cryo-SXT data, acquired under conditions that preserve cellular architecture in a near-native state, indicated that ORF3a was not associated with LDs. Collectively, these findings identify ORF3aWu as a central regulator of mitochondrial architecture and lipid storage, coupling organelle dysfunction with extensive LD expansion across distinct cellular contexts.

### A single ORF3a Omicron mutation overrides the membrane remodelling phenotype of Wuhan protein

A key question was whether the highly conserved T223I substitution present in Omicron ORF3a protein (ORF3aOmi) preserves such membrane-remodelling phenotype induced by Wuhan protein (ORF3aWu). To address this question, A549 cells stably expressing ORF3aOmi (**Figure S1**) were generated and directly compared with ORF3aWu-expressing cells. In striking contrast to ORF3aWu, ORF3aOmi did not disrupt Golgi organization. Confocal microscopy showed that the interconnected tubular Golgi network was largely preserved, rather than fragmenting into disperse puncta as observed in ORF3aWu-expressing cells **(Figure 4A-C)**. Quantitative analysis further demonstrated that ORF3aOmi occupied a Golgi surface area comparable to that of ORF3aWu (**Figure S2D, E**), indicating that the T223I substitution does not impair Golgi targeting but largely abolish the ability of ORF3a to disrupt Golgi cisternal architecture.

**Figure 4.**
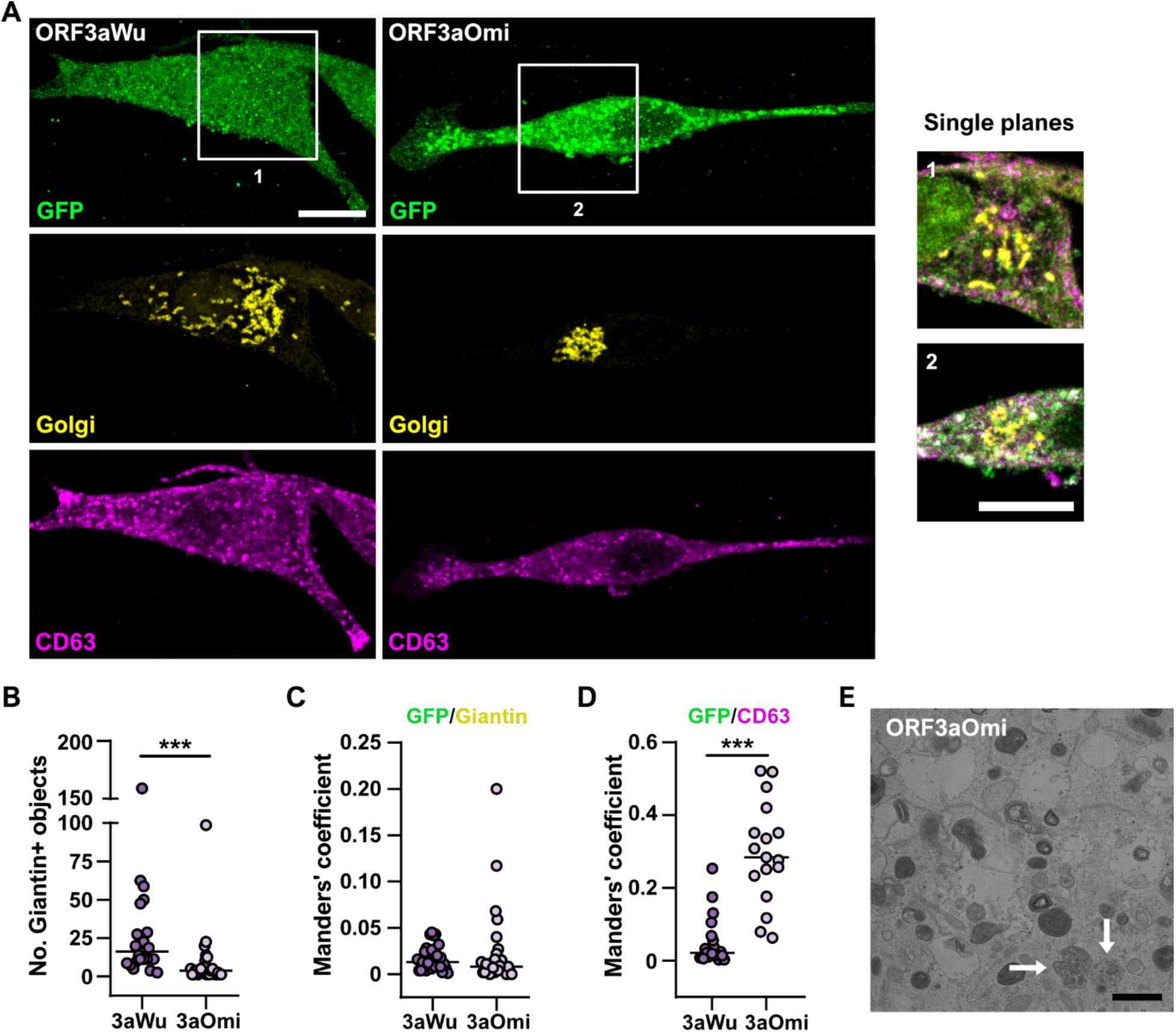
ORF3a Omicron co-localizes with multivesicular bodies (MVB) and preserves the Golgi. **(A)** Maximal projections from confocal stacks showing GFP (green), Golgi (giantin, yellow) and MVB (CD63, magenta) in A459 cells expressing ORF3aWu-GFP and ORF3aOmi- GFP. Insets 1 and 2 are shown in right panels as magnified, single plane images. Scale bar: 10 μm. **(B)** Quantification of Golgi particles (number of Giantin+ objects) in ORF3aWu (n = 28) and ORF3aOmi (n = 27) cells. **(C)** Manders’ coefficient for GFP co-localization with Giantin (ORF3aWu, n = 26; ORF3aOmi, n = 27) and **(D)** CD63 (ORF3aWu, n = 21; ORF3aOmi, n = 17) in control and ORF3aWu cells. Graphs, median. **(E)** TEM image of a 70 nm section of ORF3aOmi-transduced A549 cells. White arrows indicate dense vesicles. Scale bar: 1 µm. Statistical significance is given as: ***p < 0.001.

Instead, ORF3aOmi showed partial colocalization with CD63-positive compartments **(Figure 4A, D)**, consistent with localization to MVBs and late endosomes. TEM imaging supported these observations by revealing MVB-like structures in ORF3aOmi-expressing cells **(Figure 4E)**. These findings suggested that T223I substitution shifted ORF3a activity from Golgi-associated membrane remodelling towards the endolysosomal pathway while preserving Golgi integrity. Because Golgi architecture and endolysosomal trafficking are intimately linked to intracellular lipid trafficking, autophagic flux and membrane recycling, this redistribution would be expected to profoundly alter cellular lipid homeostasis. Consistent with this interpretation, ORF3aOmi failed to produce the metabolic phenotype induced by Wuhan protein. Unlike ORF3aWu, ORF3aOmi did not significantly increase the proportion of cells with low mitochondrial membrane potential or promote LD accumulation, as determined by MitoView and LipidTOX staining, respectively (**Figure S3B, Figure 5A, B**). These observations agree with previous work showing that residues S171, W193, and L219 within the ORF3a ‘cytosolic domain’ are critical determinants of LD accumulation and autophagy inhibition^22^. Although these residues are conserved in ORF3aOmi, the Omicron protein may contain substitutions at K67N, T89I and, most notably, T223I. Previous studies demonstrated that the T223I substitution alone (as in the ORF3aOmi under study) is sufficient to weaken the interaction between ORF3a and the HOPS complex subunit VPS39, reduce LD accumulation and attenuate viral replication^22^, strongly suggesting that this single amino acid substitution is largely responsible for the phenotypic differences observed here.

**Figure 5.**
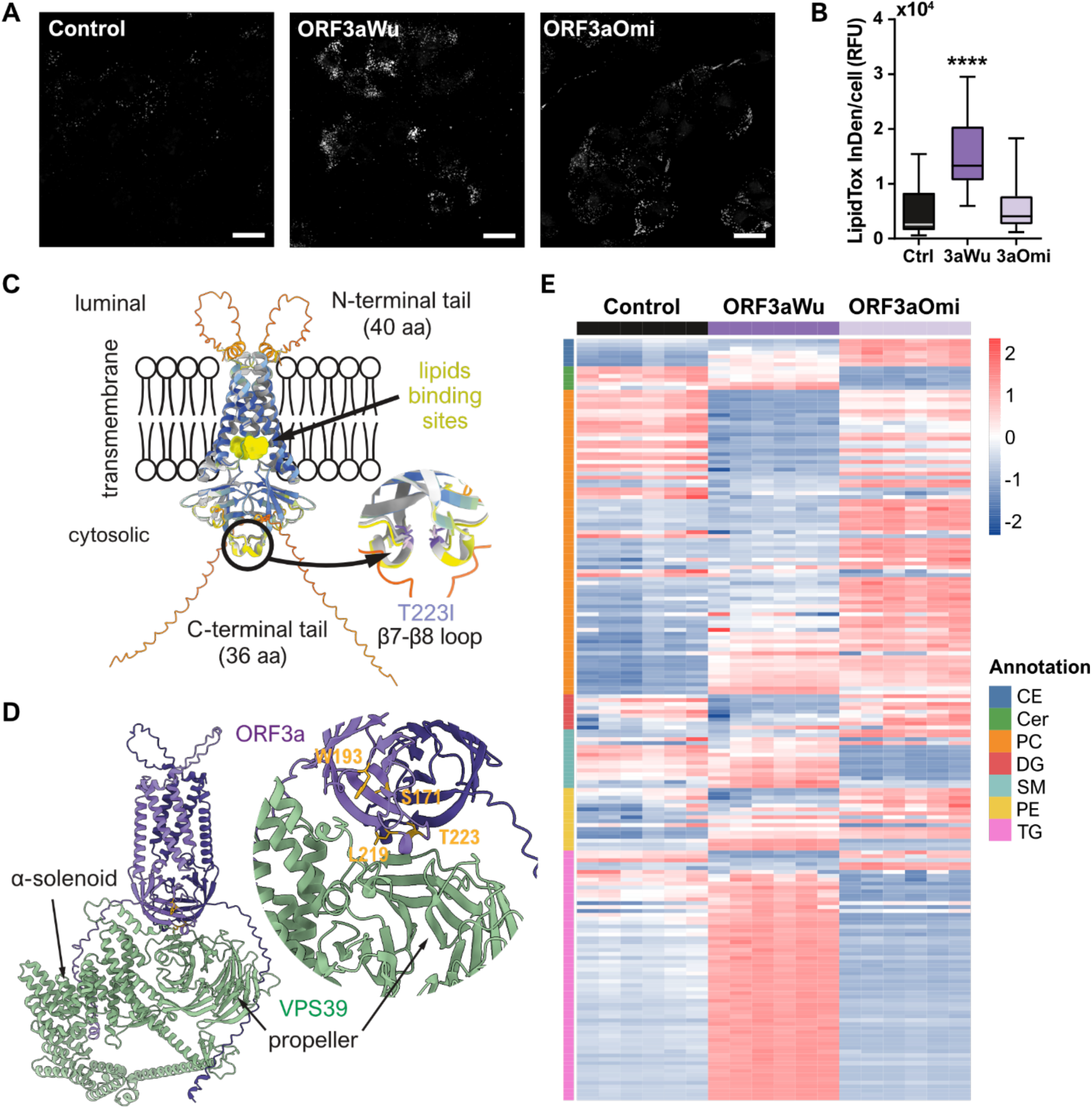
ORF3a Wuhan but not Omicron induces LD accumulation. **(A)** Representative confocal images of A549 cells stained with LipidTox. LD appear white over the black background (objective: 63x, scale bar: 25 µm). **(B)** Box and whisker plot showing the distribution of the integrated density of LipidTox fluorescence per cell in A549 control (n = 33), ORF3aWu (n = 36) and ORF3aOmi (n = 40) cells. **(C)** AlphaFold3 model of the ORF3aT223I dimer (cartoon representation) coloured by predicted local distance difference test (pLDDT) confidence values and showing the luminal and cytosolic flexible tails. The experimentally determined structure (PDB 7kjr) is superimposed to highlight the lipid-binding sites (yellow). The inset shows the superposition of both structures, revealing no detectable conformational changes in the β7–β8 loop due to the T223I substitution. **(D)** AlphaFold3 model of the ORF3a–VPS39 complex (cartoon representation). ORF3a (purple) interacts with the β-propeller domain of VPS39 (green). The inset shows the predicted binding interface, highlighting ORF3a residues S171, W193, L219 and T223 (orange), which have been implicated in the interaction with VPS39. **(E)** Heatmap showing altered lipids obtained in positive ionization mode in control, ORF3aWu and ORF3aOmi A549 cells, sorted by lipid families: cholesteryl esters (CE), ceramides (Cer), phosphatidylcholines (PC), diacylglycerols (DG), sphingomyelins (SM), phosphatidyl-etanolamines (PE) and triacylglycerols (TG). Color key corresponds to row-normalized values of lipid concentration. Statistical significance is as follows: ****p < 0.0001.

To investigate the structural consequences of the T223I substitution, we compared the five top-ranked AlphaFold3 predictions of ORF3aWu and ORF3aT223I. **Figure S5A** shows the comparison of experimental cryo-EM structure of SARS-CoV-2 ORF3a (PDB 7kjr) with the highest-ranked AlphaFold3 predictions of ORF3aWu and ORF3aT223I. All predicted structures showed excellent agreement with the experimental cryo-EM structure of ORF3a (PDB 7kjr), with Cα RMSDs of 0.587 – 0.631 Å and 0.463 – 0.550 Å for ORF3aWu and ORF3aT223I, respectively (**Figure S5A**). No significant local conformational differences were detected around residue 223 in any of the highest-ranked models (**Figure 5C**), although confidence in this region was low **(Figure S5A)**. Nevertheless, the corresponding region in ORF3aWu models closely matches the experimental structure. Both variants displayed comparable overall confidence scores (pTM = 0.62 and 0.66 for ORF3aWu and ORF3aT223I, respectively). Notably, despite the low confidence assigned to the intrinsically disordered regions, the top-ranked models consistently predicted a similar conformation for the 40-residues N-terminal “luminal” region, particularly for ORF3aWu, whereas substantially greater variability was observed for the flexible 36-residues ‘cytosolic’ C-terminal tail (**Figure S5A**). The consistency among independent predictions for the N-terminal regions raised the possibility that the T223I substitution was associated with a N-terminal “luminal” domain reorientation, despite the absence of detectable local structural changes at the mutation site (**Figure 5C**). If confirmed experimentally, such a rearrangement could provide a mechanism whereby T223I exert long-range allosteric effects rather than acting through local structural perturbations. In contrast, conformational heterogeneity observed among the highest-ranked models for the intrinsically flexible ‘cytosolic’ C-terminal tail precluded unambiguous attribution of structural differences in this region to the T223I substitution.

Given the previously reported interaction between ORF3a and VPS39, mediated by the C-terminal ‘cytosolic domain’ of ORF3a, we used AlphaFold3 to explore the structural basis of this interaction and, in particular, to determine whether ORF3a might compete with Rab7 for VPS39 binding. Recent AlphaFold modelling of the VPS39-Rab7 complex positioned Rab7 on the N-terminal α-solenoil region of VPS39, providing a structural framework for HOPS recruitment to late endosomal membranes^38^. We therefore asked whether ORF3a recognizes the same interface or instead engaged VPS39 through a distinct molecular mechanism. AlphaFold3 modelling consistently predicted ORF3a binding to VPS39 β-propeller domain rather than to Rab7-binding α-solenoid region (**Figure S5B**). Moreover, several of the predicted interaction interfaces were consistent with experimental evidence indicating that ORF3a β2-β3 loop is the major contribution to VPS39 binding (**Figure 5D**). Although these structural models required experimental validation, they suggested that ORF3a unlikely inhibits HOPS function simply by competing with Rab7 for VPS39 binding. Instead, ORF3a may remodel VPS39 or alter its orientation within the HOPS complex, thereby perturbing membrane tethering and redirecting intracellular membrane trafficking (**Figure S5C**).

Such a mechanism is consistent with the increasingly diverse functions attributed to VPS39 beyond its canonical role in membrane fusion. In addition to its established role in membrane trafficking, VPS39 has been implicated in several mitochondrial and lysosomal processes. VPS39 facilitates phosphatidylethanolamine (PE) transport to mitochondria ^39^, is recruited to mitochondria in response to sphingolipid signaling ^40^, and contributes to the formation of vacuole–mitochondria contact sites (vCLAMPs) through interaction with the outer mitochondrial membrane protein TOM40^41^. Furthermore, recent evidence has identified VPS39 as a regulator of lysosomal cholesterol export, acting through modulation of NPC2 trafficking and bis(monoacylglycerol)phosphate (BMP) metabolism^42^. Interestingly, ORF3a has also been reported to contain multiple lipid-binding pockets^5,6^ and molecular dynamic simulations have suggested that it may facilitate lipid transportation across the membrane^43,44^. Together, these observations support the emerging view that VPS39 functions not only as a membrane-tethering factor but also as a central coordinator of intracellular lipid homeostasis, raising the possibility that ORF3a may exploit this pathway to remodel host membrane composition and lipid trafficking. In light of these findings, and considering the marked differences in cellular remodelling induced by ORF3aWu and ORF3aOmi, we propose that disruption of the ORF3a-VPS39 axis by the T223I substitution largely uncouples the extensive membrane remodelling and metabolic rewiring induced by the Wuhan ORF3a protein.

LDs constitute the primary intracellular storage organelles for neutral lipids, containing the majority (50 - 90 %) of cellular triacylglycerols (TGs) together with variable amounts of cholesteryl esters (CEs)^45^. Therefore, we next investigated whether ORF3aWu and ORF3aOmi distinct phenotypes were accompanied by specific lipidomic changes. Lipidomic analysis revealed a selective accumulation of TGs species in ORF3aWu-expressing cells relative to both control and ORF3aOmi cells **(Figure 5E and S6)**, whereas total CE levels remained essentially unchanged. Interestingly, ORF3aOmi exhibited a differential pattern from control and ORF3aWu cells in cholesteryl esters (CE), ceramides (Cer), phosphatidylcholines (PC), sphingomyelins (SM) and phosphatidyl-etanolamines (PE) **(Figure 5E)**. Collectively, these data demonstrated that TG accumulation as the primary driver of LD expansion in ORF3aWu-expressing cells and further indicating that T223I substitution largely suppressed Wuhan ORF3a protein metabolic rewiring.

### ORF3aWu upregulates Triacylglycerol biosynthetic pathway

To define the metabolic basis of this phenotype, transcriptional changes in lipid biosynthesis pathways were analysed. RNA-seq revealed selective upregulation of enzymes involved in TG synthesis in ORF3aWu **(Figure S7)**. Specifically, considering LD biogenesis pathway, 1-acylglycerol-3-phosphate acyltransferase AGPAT4, and phosphatidic acid phosphatase LPIN2 increased significantly, whereas other isoforms remained unchanged or decreased **(Figures 6A and S7)**. Notably, both diacylglycerol acyltransferases DGAT1 and DGAT2, which catalyse the final step of TG synthesis, showed significant upregulation. In contrast, sterol O-acyltransferase SOAT1 and SOAT2, which mediate CE synthesis, did not change, consistent with the absence of CE accumulation in our lipidomic results **(Figures 5E and S6)**. These data linked ORF3aWu expression to specific activation of TG biosynthetic pathway.

**Figure 6.**
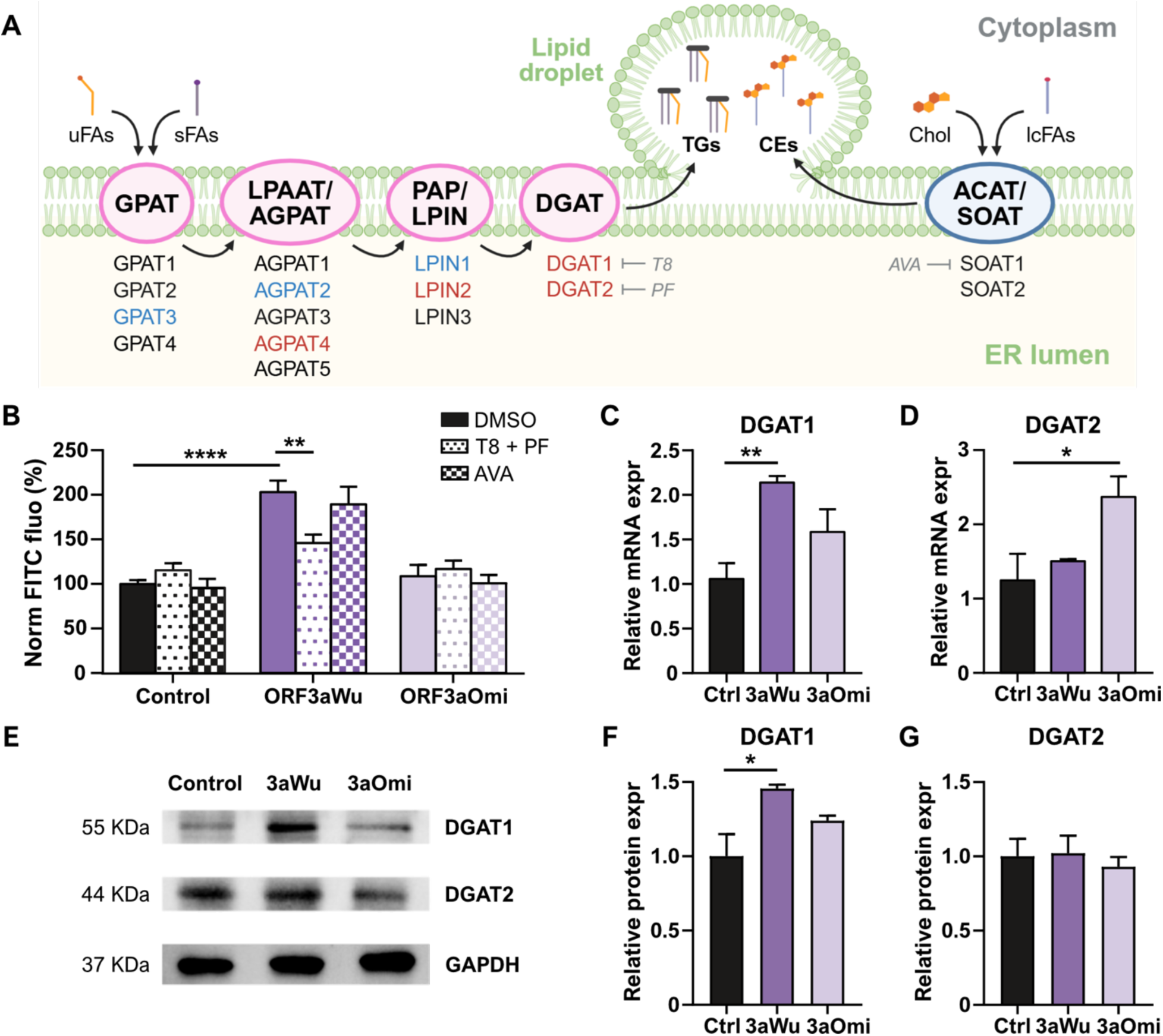
ORF3aWu drives LD accumulation by increasing DGAT1 expression in the TG synthesis pathway. **(A)** Schematic view of LDs’ biogenesis in the ER membrane via TGs (pink) and CEs (blue) metabolism. Enzymes (rounded) and their isoforms are shown; blue and red colours correspond to downregulated and upregulated genes respectively in ORF3aWu versus control A549 cells in transcriptomic analysis. Selective inhibitors of some isoforms are shown in grey (T8: T863, PF: PF-06424439, AVA: Avasimibe). sFAs: saturated fatty acids, uFAs: unsaturated fatty acids, Chol: cholesterol, lcFAs: long chain fatty acids. Figure created with Biorender.com. **(B)** Percentage of normalized FITC fluorescence in Bodipy-labelled control, ORF3aWu and ORF3aOmi A549 cells following overnight treatment with 1% DMSO, 10 μM AVA, or an equimolar combination of T8 and PF at 5 μM each. **(C)** Relative expression of DGAT1 and **(D)** DGAT2 genes validated by qPCR. **(E - G)** Western Blot and relative quantification of DGAT1 and DGAT2 proteins. Statistical significance is given as: *p < 0.05, **p < 0.01, ****p < 0.0001. In all cases data are represented as mean ± SEM (n = 3).

Subsequently, TG synthesis to LD accumulation was assessed by flow cytometry quantifying neutral lipids using BODIPY 493/503 on control, ORF3aWu-and ORF3aOmi-expressing cells in the presence of different inhibitors. Because DGAT1 and DGAT2 enzymes can compensate for each other^46–48^, we inhibited both simultaneously using T863 and PF-06424439 (T8 and PF). As control, we also inhibited SOAT1, since this protein is widely expressed systemically^49^, using avasimibe (AVA). Consistent with immunofluorescence results **(Figure 5A, B)**, ORF3aWu cells displayed a significantly higher neutral lipid content under basal conditions **(Figure 6B)**. DGAT inhibition (T8 + PF) significantly reduced neutral lipid levels in ORF3aWu cells, whereas AVA had no effect. In contrast, ORF3aOmi cells did not exhibit a significant increase in neutral lipid content relative to control cells, and none of the treatments produced a significant effect **(Figure 6B)**, suggesting that our conditions were insufficient to affect basal neutral lipids levels in those cells. Instead, DGAT inhibition in ORF3aWu cells was sufficient to reduce neutral lipid accumulation, likely due to the overexpressed levels suggested by RNA-seq. AVA showed no effect, consistent with SOAT1 absence of overexpression (**Figures 6A and S7)**.

Since DGAT1 and DGAT2 are distinct family enzymes^50,51^, we further analysed the putative selectivity of ORF3a for one isoform. In ORF3aWu cells, qPCR showed a ∼ 2-fold increase in DGAT1 expression, whereas DGAT2 levels remained unchanged **(Figure 6C, D)**. Western blot analysis confirmed selective upregulation of DGAT1 protein level **(Figure 6E, F)**. In ORF3aOmi cells, DGAT1 expression increased slightly but did not reach statistical significance and DGAT2 protein levels remained unchanged despite increased mRNA **(Figure 6C-E)**. Taken together, our results showed that single T223I substitution in ORF3a rewires protein function. Whereas the original strain from Wuhan disrupts Golgi architecture, drives DGAT1-dependent TGs synthesis and LDs expansion, the substitution of a polar Thr by a hydrophobic Ile in Omicron mutant preserves Golgi integrity and fails to induce lipid accumulation. Further experiments will determine whether ORF3a interaction with DGAT1 is direct or whether it is due to an alteration of a metabolic pathway.

Our results suggested that ORF3a-VPS39 interaction may function as a lipid-interacting membrane remodelling protein rather than solely as an autophagy inhibitor. Autophagosome formation depends on ATG2-driven lipid transfer from the ER to the growing phagophore^52^. Beyond its role in autophagy, ATG2 has been reported to regulate LD biogenesis by modulating DGAT activity^53^. Consequently, ORF3a–VPS39 interaction could impair the normal function and distribution of ATG2, redirecting lipid flux toward LD expansion at the expense of membrane supply for phagophore elongation and autophagosome biogenesis. Also, cryo-EM structures of ORF3a within nanodiscs revealed discreate lipid-binding sites and lipid-filled membrane openings, supporting direct interaction with lipids and raising the possibility that ORF3a actively reorganize lipid-rich membrane domains^5,6^. Consistently, molecular dynamics simulations showed that cholesterol and phospholipids modulate ORF3a conformational dynamics and membrane-associated organization^54^. In this context, ORF3a-VPS39 interaction may redirect VPS39-associated lipid trafficking pathways away from their physiological role in organelle homeostasis toward the generation of virus-associated membrane compartments, mitochondria remodelling, and DGAT1-dependent LD biogenesis. Whether all ORF3aWu effects reported here were driven by specific interactions with VPS39 that subsequently altered DGAT1 function, or instead resulted from the extensive cellular remodelling induced by the viral protein that ultimately gave rise to such functional alterations, remains an open question. Notably, these mechanisms are not mutually exclusive and may act synergistically. Interestingly, there are striking conceptual parallels between ORF3a and African Swine Fever Virus CP204L, which also binds VPS39 preventing its recruitment to lysosomal membranes and rewires endolysosomal trafficking^55^, suggesting that such interactions may represent a common viral strategy for remodelling the cellular landscape in a manner that favours viral fitness.

## CONCLUSIONS

Our findings establish SARS-CoV-2 ORF3a as a multifunctional regulator of host-cell architecture coupling membrane remodelling with metabolic reprogramming. We showed that ORF3a from the ancestral Wuhan strain disrupted Golgi organization, promoted the formation of specialized Golgi-associated multivesicular membrane assemblies, remodelled mitochondria ultrastructure, and drove DGAT1-dependent TG synthesis as well as LD biogenesis. In contrast, our study uncovered previously unrecognized effects of the Omicron-associated T223I substitution, demonstrating that it largely abrogated these activities, preserving Golgi integrity, redirecting ORF3a to CD63-positive endolysosomal compartments and preventing mitochondrial dysfunction as well as lipid accumulation. Structural modelling further suggested that these functional changes arose from long-range allosteric effects of the T223I substitution rather than mainly local structural perturbations at the mutation site. Although our reductionist cellular models did not fully recapitulate the complexity of SARS-CoV-2 infection, it provided a robust framework for defining ORF3a-specific functions and suggested that variant-dependent differences in ORF3a activity may contribute to distinct cellular response associated with viral evolution. Together, these results provided a mechanistic link between SARS-CoV-2 genetic diversification and host-cell remodelling, identifying DGAT1-mediated lipid metabolism as a central downstream effector of ORF3a, highlight this pathway as a potential therapeutic target.

## METHODS

### Cell culture, lentivirus production and transduction

A549 alveolar basal epithelial cells line (ATCC CRM-CCL-185; RRID: CVCL_0023) and THP-1 monocytic cell line (ATCC TIB-202; RRID: CVCL_0006) were cultured in Dulbecco’s Modified Eagle Medium (DMEM) (Gibco, #41966029) and Roswell Park Memorial Institute (RPMI) (Gibco, #11875093) respectively, supplemented with 10% (v/v) heat-inactivated foetal bovine serum (FBS) (Gibco, #1027016) and 1% Penicillin-Streptomycin (100U/ml) (Gibco, #15070063) at 37 °C in a 5% CO_2_, 90% humidity atmosphere. All cultured cells were performed under these conditions except otherwise stated. ORF3a coding sequences (codon optimized for mammalian expression) were cloned into pLVXEF1a-IRES-Puro Cloning and Expression Lentivector (Clontech, Takara, #631253) to generate pseudotyped lentiviral particles encoding the ORF3a accessory protein of SARS-CoV-2 (Wuhan or Omicron isolates) at the Centro Nacional de Investigaciones Cardiovasculares Viral Vector Unit (Madrid, Spain) as previously described^56,57^. ORF3a accessory protein was either C-terminally fused with 2x StrepTag (ST) or enhanced GFP (GFP). A549 cells were incubated with ORF3a-encoding lentiviral particles at a multiplicity of infection (MOI) of 30 for 24 h, followed by 2 μg/mL puromycin (Thermo Fisher, #A1113803) treatment to select successfully transduced cells. THP-1 cells were transduced with the lentiviral particles at a MOI of 150 for 48 h, after which a second transduction was performed under identical conditions. Following a 24 h incubation, 2 μg/mL puromycin was added for cell selection. For both cell lines, control cells that expressed either ST or GFP alone were also generated. Polybrene (Merck, #TR-1003-G) was used at a final concentration of 5-8 μg/mL to improve transduction efficacy.

Viral transcripts were confirmed by qPCR^35,57^, using the primers listed in **Tables S1**. Flow cytometry was also used to confirm protein expression in ORF3a-GFP-expressing cells for which, 2 x 10^5^ cells were harvested, washed with PBS and resuspended in PBS. GFP signal was analysed using a CytoFLEX flow cytometer (Beckman Coulter) and FlowJo v10 software (BD Biosciences).

### Transmission electron microscopy (TEM) of 70 nm cells sections

A549 cells were seeded at 2.5 x 10^5^ cells per well in 6-well plates and were cultured for 24 h. Cell monolayers were *in situ* fixed for 1 h at room temperature (RT) with 3% glutaraldehyde (EM Grade, Ted Pella INC) in PBS. Subsequently, cells were treated at 4 °C for 1 h with 1% osmium tetroxide (EM Grade, Ted Pella INC) and 0.8% potassium ferricyanide (Electron Microscopy Sciences). After dehydration in a 30 % to 100 % ethanol gradient, cells were embedded in a gradient of ethanol/LX 112 epoxy resin up to 100 % epoxy resin (Ladd Research). Samples were polymerized at 60 °C for 2 days and 70 nm sections were obtained with a Leica EM UC7 ultramicrotome (Leica Microsystems GmbH), placed on Formavar/Carbon 100 mesh copper grids (Electron Microscopy Sciences, #FCF-100-CU-50) and stained 30 min with 5 % Uranyl Acetate and 3 min with 0.4% Reynold’s Lead Citrate (Sigma). Grids were observed using a Thermo Fisher TALOS L120C microscope operated at 120 kV and images were taken under low dose conditions with a Thermo Fisher CETA-F camera, using Velox and MAPS software (Thermo Fisher).

### Cryo-electron tomography of 250 nm cells sections

Same samples prepared for 70 nm sections were used to get ∼ 250 nm sections, which were placed onto Cu-Quantifoil® S7/1 grids. Grids were pre-irradiated on a JEOL JEM-1400 Flash TEM operating at 120 kV using SerialEM (1,2), and subsequently loaded into a Thermo Fisher Scientific Talos Arctica operating at 200 kV for cryo-ET data collection. Tilt series were acquired using Thermo Scientific Tomography 5 software (v5.21; Thermo Fisher Scientific, USA) with beam-image shift compensation for multiple tilt series per target area. Data were collected at a nominal magnification of 22,000 x (pixel size 0.465 nm/pixel) over an angular range of - 60° to + 60° in 1° increments, to a total accumulated dose of 140 e⁻/U. Tomograms were reconstructed on-the-fly using the Tomo Live pipeline integrated within the Thermo Scientific CryoFlow–CryoTomo software environment (Thermo Fisher Scientific, USA).

### Immunofluorescence and co-localization assays

A549 cells were seeded at 6 x 10^4^ per condition on precision cover glasses thickness no. 1.5H (Marienfeld, Germany) and were cultured for 24 h. Cells were then fixed with PHEM solution (30 mM PIPES, 20 mM HEPES, 2 mM EGTA, 1 mM MgCl_2_, pH 6.9) containing 4% paraformaldehyde (PFA) and 0.12 M sucrose for 15 min at RT. Cells were then permeabilized in PHEM containing 0.2 % TX-100, 0.5 % PFA and 0.12 M sucrose for 5 min at RT, and blocked with PHEM containing 0.5 % TX-100, 100 μg/mL γ-globulin, 3 % bovine serum albumin (BSA) and 0.2 % azide for 30 min at RT. Cells were sequentially stained with the primary and secondary antibodies shown in **Table S2**. Samples were mounted on Prolong Gold (Invitrogen P36930) and series of fluorescence and brightfield images were captured using a Leica STELLARIS navigator confocal microscope equipped with a pulsed WLL (range, 470-670 nm) and a HC PL Apo CS2 100×/1.4 oil objective (Leica Microsystems). Tau separation mode was used to acquire fluorescence corresponding to Alexa 546 lifetime. For analysis, Z-stacks were processed with Image J software. Cells were segmented and subjected to JACoP and 3D object counter v1.5.1 plugins, to find Manders’ coefficient for colocalization (GFP to CD63 and to Giantin) and to quantify the number of particles, surface and volume occupied by the Golgi, respectively^58^.

### Cryo-soft X-ray tomography (cryo-SXT) and cryo-3D-structured illumination microscopy (cryo3DSIM)

#### Grids preparation

A549 cells were seeded at 2 x 10^5^ cells/mL on top of Quantifoil® R2/2 Au G200F1 grids (Quantifoil Micro Tools GmbH, #N1-C16nAuG1-01), which were previously sterilized by UV irradiation at RT for 3 h. THP-1 cells were seeded at 2 x 10^5^ cells/mL and differentiated into macrophages by treatment with 25 ng/mL phorbol 12-myristate 13-acetate (PMA) for 48 h followed by extra 24 h incubation without PMA onto Quantifoil® R2/2 Au G200F1 grids, which were coated with 0.01% poly-L-lysine (Sigma Aldrich, #P4707) solution for 30 min at 37 °C upon UV treatment. Cells were cultured until they reached ∼ 70 % confluence and 300 nM LysoTracker Red DND-99 (Thermo Fisher, #L7528) was added for fluorescent labelling of acidic vesicles for 10 min, followed by the addition of 100 nm colloidal gold nanoparticles (BBI Solutions, #EM.GC100/7). Cells were immediately vitrified in liquid ethane in a Vitrobot Mark IV (Thermo Fisher) and stored in liquid nitrogen. The grids were screened for ice thickness and fluorescent signal on an Axio Scope A.1 (Zeiss) widefield epifluorescence light microscope and selected samples were transferred to the cryo3DSIM.

#### Cryogenic 3D structured illumination microscopy

A custom-built cryogenic 3D structured illumination microscope (cryo3DSIM) was developed at the ALBA Synchrotron and installed adjacent to the MISTRAL beamline to enable an integrated cryo-correlative fluorescence and soft X-ray tomography workflow. The microscope was designed to image vitrified specimens on standard 3 mm gold Quantifoil TEM grids, allowing direct transfer to MISTRAL cryo-SXT end station.

Selected grids with green signal coming from ORF3a-GFP protein and red signal from LysoTracker, were imaged. Samples were maintained below the vitreous ice transition temperature using a Linkam CMS196 cryostage. Fluorescence excitation was provided by 488 nm (LuxX488-200, Omicron) continuous-wave lasers at 40 mW for 50-200 ms depending on the efficient of the fluorescence. Structured illumination patterns were generated using a spatial light modulator, and three-dimensional image stacks were acquired with a piezoelectric z-stage (P-528.ZCD, Physik Instrumente). A 100X 0.9 NA objective lens was used, achieving a lateral resolution of roughly 210 nm full width at half maximum (FWHM) at an emission wavelength of 488 nm and 590 nm for GFP and lysotracker respectively. For each sample-type, three to six cells, preferably from different grids, were imaged. The system supports automated fluorescence mosaics for grid navigation and high-resolution 3D imaging of selected cells for subsequent correlation with cryo-SXT.

#### Cryo-SXT

The selected frozen grids were transferred to Mistral (ALBA-Light Source)^33,59^ beamline in ALBA synchrotron under cryogenic conditions. We used a photon energy (520 eV) within the water window to take advantage of the high natural absorption contrast of the biological material to acquire X-ray tomography data sets with the conditions described previously^60^. A tilt series was acquired for each cell area using an angular step of 1° on a ±70° range with a Fresnel Zone plate (FZP) of 40 nm outermost zone width and an effective pixel size of 13 nm. The image stacks were pre-processed to normalize and correct the intensity distribution delivered to the sample by the capillary condenser lens. Each set of projections (termed tilt series) was aligned to a common sample tilt axis using a custom-written automated patch-tracking algorithm^61^. The aligned tilt series were reconstructed into three-dimensional tomograms using the tomo3d software (SIRT algorithm with 50 iterations)^62^. To enhance the signal-to-noise levels, we used TOMOEED^63^. The spatial resolution of the final 3D reconstructions was determined by the Fourier Shell Correlation (FSC) even/odd method^64^, yielding a half-pitch resolution of 27.7 nm at 0.25 FSCe/o.

The resulting volumes were visualized and segmented using Amira 3D software (ThermoFisher Scientific) and ChimeraX^65^. For both A549 and THP-1 PMA-differentiated cells, we selected eight tomograms per condition from independent cells and segmented the volumes corresponding to the LD and the mitochondria. They were divided in two categories, according to their electron density and cristae’s appearance: if the mitochondria were highly electrodense and/or their cristae were deteriorated, they were considered as altered (“bad” group), otherwise they were classified as normal (“good” group). To perform the quantitative analysis of the morphology of these organelles, we used the Mitochondria Analyzer plugin for ImageJ software.

#### Correlative cryo-SXT / cryo3DSIM imaging

The cryo-SIM and cryo-SXT datasets were registered using the multidimensional correlation software eC-CLEM^66^. An initial coarse alignment was performed by calculating a two-dimensional rigid transformation between the SIM images and the X-ray tomograms. This transformation was obtained through a sequence of independent 2D registrations: (i) cryo-SIM fluorescence data were aligned to the corresponding brightfield image stacks, (ii) the brightfield images were registered to the 2D X-ray mosaics, and (iii) the X-ray mosaics were aligned with the positions of the X-ray tomograms. The resulting 2D transformation was subsequently applied to bring the 3D cryo-SIM and cryo-SXT regions of interest into approximate correspondence, providing the starting point for the full three-dimensional registration. Alignment relied on intrinsic features visible in both imaging modalities, including grid landmarks (grid bars, holes in the carbon support film and sample imperfections) together with cellular structures exhibiting sufficient contrast in fluorescence and X-ray microscopy, such as fluorescent endosomes. Features present immediately before vitrification were also used when appropriate. The final registration step consisted of a rigid three-dimensional alignment of the chromatic aberration-corrected cryo-SIM volumes with the corresponding cryo-SXT tomograms using the 3D registration mode implemented in eC-CLEM^66^. Only rigid transformations were applied during the registration process, allowing translation, rotation and isotropic scaling, while no non-rigid deformations were introduced. Colocalization analysis was performed on aligned fluorescence image stacks using the Coloc2 plugin implemented in Fiji/ImageJ^67^. Pearson’s correlation coefficient (PCC), thresholded Manders’ coefficients^68^, and Costes randomization test^69^ were used to quantify and assess the significance of colocalization. The correlated cryo-SXT and cryo3DSIM volumes were visualized using UCSF ChimeraX^65^.

### Mitochondrial activity

Flow cytometry was used to visualize cell populations according to their mitochondria activity as previously described.^4,35^

### Lipid droplets staining and analysis

A549 cells were seeded at 4 x 10^4^ per well were seeded on top of #1 glass coverslips in 24-well plates and cultured for 24 h. Cells were fixed with 4 % PFA in DMEM for 5 min at 37 °C and stained with a 1:1000 dilution of LipidTOX Green (Invitrogen, #H34475) for 2h at RT. Coverslips were mounted on FluorSave^TM^ Reagent (Merck, #345789) and images were acquired with a confocal laser microscope Leica TCS SP8 STED 3X.

For neutral lipid accumulation assay, A549 cells were seeded at 8 x 10^4^ per well in 24-well plates and incubated for 6 h at 37 °C. Cells were then treated with either 1 % dimethyl sulfoxide (DMSO) (ITW Reagents, #A3672), 10 μM Avasimibe (MedChemExpress, #HY-13215), or a combination of 5 μM T863 (MedChemExpress, #HY-32219) and 5 μM PF-06424439 (MedChemExpress, #HY-108341). Following overnight incubation at 37 °C with 5 % CO_2_, cells were labelled with 10 μM BODIPY 493/503 (MedChemExpress, #HY-W090090) for 15 min at RT in the dark, and resuspended in 200 μL of PBS buffer containing 0.5 % BSA, 2 mM Ethylenediaminetetraacetic acid (EDTA) and 1 μg/mL 4′,6-diamidino-2-phenylindole (DAPI) (ThermoFisher Scientific, #62248). Samples were kept on ice and darkness until data acquisition using a CytoFLEX flow cytometer (Beckman Coulter).

Mean fluorescence intensity (MFI) in the green FITC channel was measured by flow cytometry using the viable (DAPI-negative) cell population. Cells not labelled with BODIPY 793/503 were used as a negative control for green autofluorescence, and their fluorescent signal was subtracted from the analysis. Data was analysed with FlowJo v10 software (BD Biosciences).

### AlphaFold 3 complex formation prediction

The three-dimensional structures of the protein sequences were predicted using AlphaFold 3^70^. Predictions were generated using the default settings and databases available at the time of analysis. AlphaFold 3 integrates information from multiple sequence alignments (MSAs), structural templates, and diffusion-based deep-learning models to predict protein structures at atomic resolution.

### Lipidomics

A549 cells were seeded at 4 x 10^5^ per condition in 6-well plates overnight (six replicates per condition), cultured for 24 h, washed with PBS and stored at – 80 °C. Samples were processed by the Metabolomics Platform at CIC bioGUNE (Bizkaia, Spain) and analysed in an ultrahigh performance liquid chromatography system (Acquity, Waters Inc., Manchester, UK) coupled to a Time-of-Flight mass spectrometer (SYNAPT G2, Waters Inc.). The mass spectra data were acquired in positive and negative electrospray ionization modes using the MassLynx V4.2 software (Waters). Lipid families of the obtained features were identified based on the mass-to-charge ratio (*m/z*) of the metabolite and the fragmentation pattern obtained by tandem mass spectrometry (MS/MS) analysis and comparing them with in-house databases as well as publicly available databases^71–76^. Unsupervised Principal Component Analysis (PCA) was used to assess the reliability of the analytical procedure and visualize tendencies among sample groups. The R package “pheatmap” V1.0.13 was used to build a heatmap of feature concentrations ordered by lipid family and cell line.

### Transcriptomic analysis

#### Sample preparation

1.5 x 10^6^ A549 cells were collected to perform RNA isolation and sequencing as previously described^57^. Each sample was collected in quadruplicate.

### Gene sets and differential gene expression analysis

Sequencing raw data was quality controlled (error rate, GC content distribution) and filtered, removing bad quality and N-containing sequences and adaptors. Clean data were mapped with HISAT2 v.2.0.5 to reference genome GRCh38.p13, and gene expression was quantified using featureCounts v.1.5.0-p3.^77^ R software was used to evaluate the differential expression among the experimental groups (ORFs vs control). Genes with fewer than 10 counts in at least 3 samples were filtered out to improve statistical power and reduce noise. DESeq2 v.1.46.0 was used to obtain differentially expressed genes, employing Benjamini– Hochberg correction and a significance cutoff of adjusted p-value < 0.05.^78^ Apeglm v.1.28.0 was used to shrink the resulting log-fold change values.^79^ Expression values were normalized by variance stabilizing transformation. These values were used to perform principal component analysis (PCA) plots with ggplot2 v.4.0.0 and heatmaps of gene expression with “pheatmap” V1.0.13. Functional enrichment analysis of differentially expressed genes was performed using clusterProfiler v.4.14.6, and the results were visualized with enrichplot v.1.26.6, considering significantly enriched terms with an adjusted p-value < 0.05.^80,81^

### Real time qPCR

cDNA was synthesized from 1 μg of total A549 cells RNA using the NZY First-Strand cDNA Synthesis Kit (Nzytech, #MB12502), according to manufacturer’s instructions. qPCR reactions were performed following the protocol provided with the NZYSpeedy qPCR Green Master Mix (Nzytech, #MB22402), using the primers sequences listed in **Supplementary Table 1**. Real time qPCR was carried out in LightCycler 96 (Roche) under the following conditions: 5 min at 95 °C followed by 45 cycles of 20 s at 94 °C and 30 s at 57 °C. Melting curve analyses were performed at the end, to confirm specificity of each PCR product. Relative expression results were calculated using GenEx6 Pro software (MultiD-Göteborg, Sweden), based on the obtained quantification values (Cq).

### Western Blot

A549 cells were seeded at 5 x 10^5^ in 6-well plates and incubated overnight at 37°C with 5% CO_2_. Cells were then harvested and lysed in ice-cold Pierce IP Lysis Buffer (ThermoFisher Scientific, #87787) at 4 °C. Total protein concentration was calculated following the Bradford method and 20 μg protein extracts were mixed with 5x SDS-PAGE Sample Loading Buffer (Nzytech, #MB11701), heated at 95 °C for 5 min, resolved by SDS-polyacrylamide gel electrophoresis and transferred onto a polyvinylidene difluoride (PVDF) membrane using a Mini Trans-Blot System (Bio-Rad, #1703935). Membranes were blocked for 1h with 5% BSA in Tris-buffered saline-Tween20 buffer and then incubated with the appropriate primary and secondary antibodies (**Table S2**). Protein bands were visualized by chemiluminescence using ChemiDoc Imaging Systems (Bio-Rad) and relative protein expression levels were quantified by sequential normalization to the loading control (GAPDH) using the Fiji software V2.1.

### Statistical analysis

Statistical analyses were performed using GraphPad PRISM 8. P-values were determined using either Student’s t-test or Mann-Whitney U test for pairwise comparisons, and one-way ANOVA followed by Bonferroni or Tukey’s post-hoc test for comparisons among multiple groups. Unless otherwise stated, data are shown as the mean of at least three biological replicates. Significant differences are indicated as: *, p <0.05; **, <0.01; ***, p<0.001, ****, p<0.0001.

## Supporting information

De Lucas et al

## ASSOCIATED CONTENT

## Supporting Information

“Characterization of transduced A549 and THP-1 cells; Analysis of Giantin distribution in A549 cells; Analysis of mitochondria structure and activity in A549 transduced cells; THP-1 cells cryo-SXT; AlphaFold3 structural models of ORF3a and its interaction with VPS39; Lipidomic analysis; Transcriptomic analysis; List of primers used for real time qPCR; List of antibodies used for immunofluorescence assays (IF) and Western Blot (WB)” (PDF).

“Movie: 3DB vesicle XZ tomogram from a A549-ORF3aWu-ST 250 nm section (related to Figure 1C)” (MP4).

“Movies: Cryo-correlative of 3DB (related to Figure 2A); Cryo-correlative of lysosome (related to Figure 2B); Cryo-correlative of 3DB-lysosome (related to Figure 2C)” (MPG).

“Movies: Segmented tomogram of transduced A549 expressing GFP (related to Figure 3A); Segmented tomogram of transduced A549 expressing ORF3aWu (related to Figure 3A)” (MP4),

## Author Contributions

Conceptualization: M.M and M.A.O. Investigation: All authors. Formal Analysis: A.L-R, U.M-H, M.P-B, A.J.P-B, O.E.A-D, F.J.C, N.B.N-C, M.A.O., M.M. Writing – Original draft: A.L-R, M.P-B, M.M, and M.A.O. Writing – Review and editing: A.L-R, M.P-B, M.M, and M.A.O. Funding acquisition: M.M, M.A.O and J.J.G. Supervision: M.M and M.A.O.

## Acknowledgments

This research work was funded by Ministerio de Ciencia e Innovación (PID2021-123399OB-I00 to M.M and M.A.O, PID2024-162356OB-I00 to M.M. and CNS2023-145079 to M.A.O), the European Commission – NextGenerationEU (Regulation EU 2020/2094) through CSIC’s Global Health Platform (PTI+ Salud Global) (COVID-19-117 and SGL2103015 to M.M.), Junta de Andalucía (CV20-20089 to J.J.G), and ALBA Synchrotron standard proposals 2022025607, 2023027348 and 20250340193 to M.A.O and M.M. This work was funded by the European Union-NextGenerationEU through the Recovery, Transformation and Resilience Plan of Spain, under Project ICT-2022-007864.

A.L-R. (PIPF-2022/SAL-GL-25558) and L.M-G. (PIPF-2023/SAL-GL-29969) are recipients of an Ayuda para la contratación de Personal Investigador Predoctoral en Formación (CAM). U.M-H. is supported by an Ayuda para contratos Predoctorales para la Formación de Doctores/as (PRE2024-002315, AEI). T.G.G. is recipient of a Ramón y Cajal contract (RYC2021-031614-I, MCIN/AEU/10.13039/501100011033 and NextGeneration EU/PRTR). R.F.R. is recipient of a Contrato Personal Investigador en Formación (PIF), University of Córdoba. J.M. is supported by Talento César Nombela (2024-T1SAL-GL-31373).

We acknowledge Rafael Nuñez Ramirez and Begoña Pou Alonso for the outstanding technical support at the Electron Microscopy Core facility at the CIB-CSIC. A.L-R, U.M-H, and L.M-G. are currently Ph.D. students at the Universidad Complutense de Madrid and their affiliation reflects their enrolment in their doctoral program.

## Abbreviations

3DBs: 3a dense bodies
Aps: accessory proteins
ARTs: algebraic reconstruction techniques
AT2: alveolar type 2
AVA: avasimibe
BMP: bis(monoacylglycerol)phosphate
BSA: bovine serum albumin
CEs: cholesteryl esters
Cq: quantification values
cryo-EM: cryo-electron microscopy
cryo3DSIM: cryo-3D-structured illuminated microscopy
cryo-SXT: cryo-soft X-ray tomography
DAPI: 4′,6-diamidino-2-phenylindole
DMEM: Dulbecco’s Modified Eagle Medium
DMVs: double-membrane vesicles
DMSO: dimethyl sulfoxide
E: envelope
EDTA: ethylenediaminetetraacetic acid
EM: electron microscopy
ER: endoplasmic reticulum
ET: electron tomography
FBS: fetal bovine serum
FSC: Fourier shell correlation
FWHM: full width at half maximum
FZP: Fresnel zone plate
GFP: enhanced green fluorescent protein
HOPS: Homotypic Fusion and Protein Sorting
LAC: linear absorption coefficient
LD: lipid droplet
M: membrane
MFI: mean fluorescence intensity
MOI: multiplicity of infection
MS/MS: tandem mass spectrometry
m/z: mass-to-charge ratio
MVB: multivesicular bodies
N: nucleocapsid
ORF3aOmi: ORF3a Omicron
ORF3aWu: ORF3a Wuhan
PCA: principal component analysis
PCC: Pearson’s correlation coefficient
PE: phosphatidylethanolamine
PFA: paraformaldehyde
PF: PF-06424439
pLDDT: predicted local distance difference test
PMA: phorbol 12-myristate 13-acetate
PVDF: polyvinylidene difluoride
RPMI: Roswell Park Memorial Institute medium
RT: room temperature
S: spike
ST: 2×Strep-tag
T8: T863
TGs: triacylglycerols
TEM: transmission electron microscopy
ToF MS: time-of-flight mass spectrometer
UPLC: ultrahigh performance liquid chromatography system
vCLAMPs: vacuole and mitochondria contact sites
ΔΨm: mitochondrial membrane potential.

The authors declare that they have no competing interest.

All data needed to evaluate the conclusions in the paper are present in the paper and/or the Supplementary Information

