## Supplementary material for "A single Omicron mutation reshapes ORF3a-driven host-cell remodelling": De Lucas et al: De Lucas et al, Supporting Information, 2 Aug26.pdf

[1]Centro de Investigaciones Biológicas Margarita Salas (CIB-CSIC), 28040, Madrid, Spain.

[2]Facultad de Medicina, Universidad Complutense de Madrid (UCM), Madrid, Spain.

[3]ALBA Synchrotron Light Source, Cerdanyola del Vallès, Barcelona, Spain.

[4]Immunology Service, Health Research Institute of La Princesa University Hospital (IIS-Princesa), Centro de Investigación Biomédica en Red Cardiovascular (CIBERCV), Madrid, Spain.

[5]Department of Anatomy and Physiology, Lithuanian University of Health Sciences, Kaunas, Lithuania.

[6]Institute of Microbiology and Virology, Lithuanian University of Health Sciences, Kaunas, Lithuania.

[7]Metabolomics Platform, & Exosomes Laboratory, CIC bioGUNE, Basque Research and Technology Alliance (BRTA), 48160 Derio, Bizkaia, Spain.

[8]Department of Macromolecular Structure, Centro Nacional de Biotecnología (CSIC), Madrid, Spain.

[9]Department of Genetics, Immunogenomics and Molecular Pathogenesis Group, UIC Zoonoses and Emergent Diseases ENZOEM, University of Córdoba, Córdoba, Spain.

[10]Maimónides Biomedical Research Institute of Córdoba (IMIBIC), Córdoba, Spain.

[11]IKERBASQUE, Basque Foundation for Science, 48009 Bilbao, Spain.

[12]Centro de Investigación Biomédica en Red de Enfermedades Hepáticas y Digestivas (Ciberehd), Madrid, Spain.

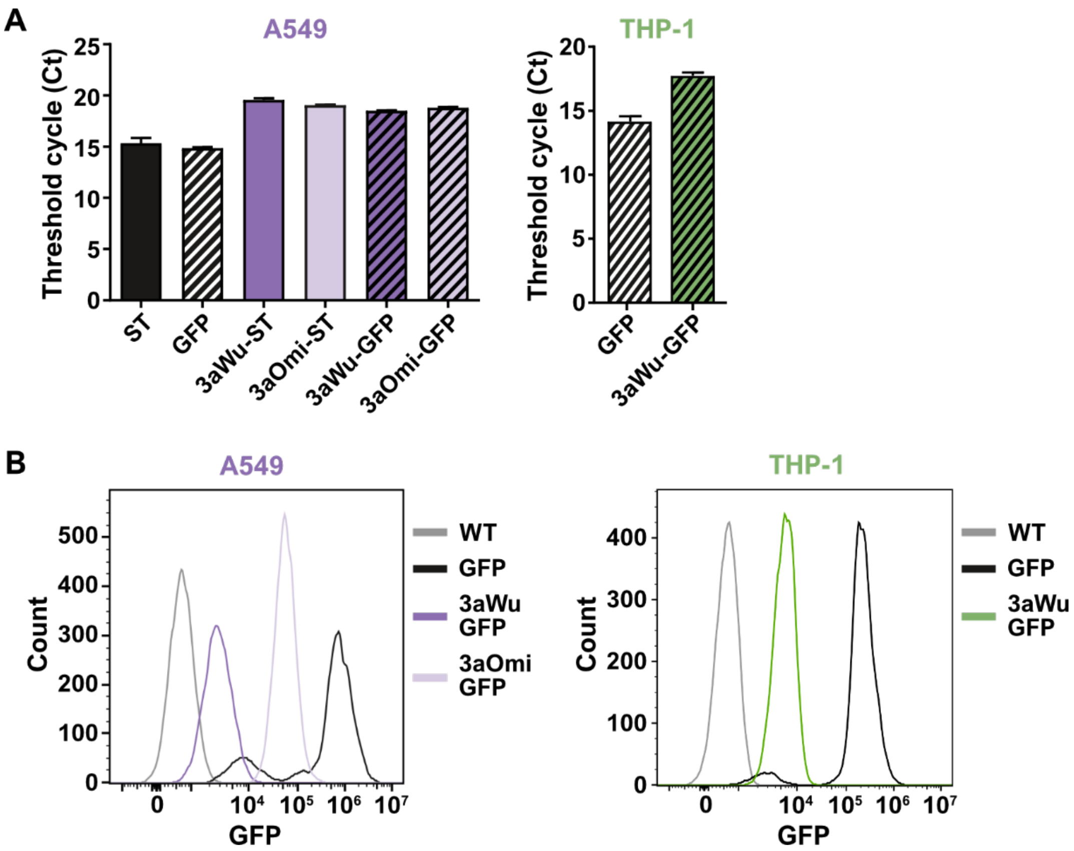


**Figure S1. Characterization of transduced A549 and THP-1 cells. (A)** Cycle threshold (Ct) values determined by real-time PCR directed to 2x Strep-Tag (ST) and GFP controls, as well as ST- and GFP-tagged ORF3aWu and ORF3aOmi constructs in transduced A549 (left) and THP-1 (right) cells. **(B)** Flow cytometry analysis of GFP expression in wild-type cells (WT, non-transduced; grey), GFP-control cells (black), and ORF3a-transduced A549 (purple) and THP-1 (green) cells. Histograms show cell counts as a function of GFP fluorescence intensity.

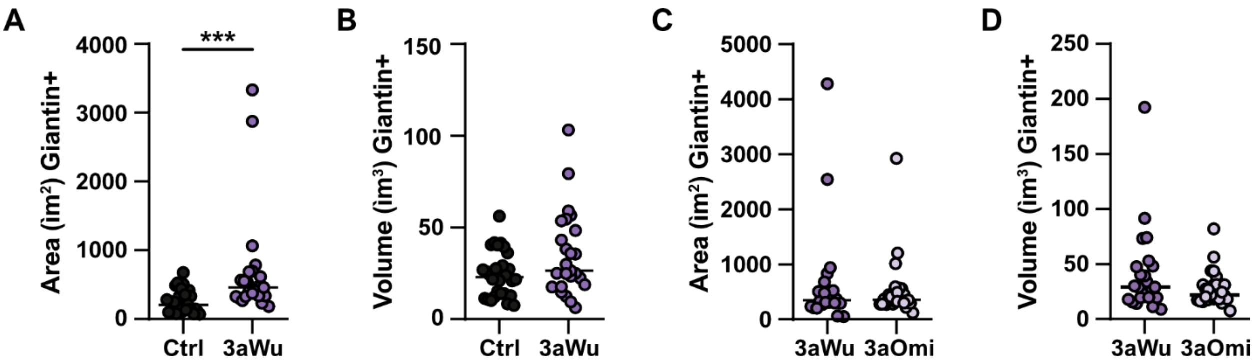


**Figure S2. Analysis of Giantin distribution in A549 cells. (A, B)** Quantification of the surface area **(A)** and volume **(B)** of Giantin-positive objects in control and ORF3aWu-expressing A549 cells. **(C, D)** Quantification of the surface area **(C)** and volume **(D)** of Giantin-positive objects ORF3aWu- and ORF3aOmi-expressing A549 cells. Statistical significance is given as: ***$p < 0.001$.

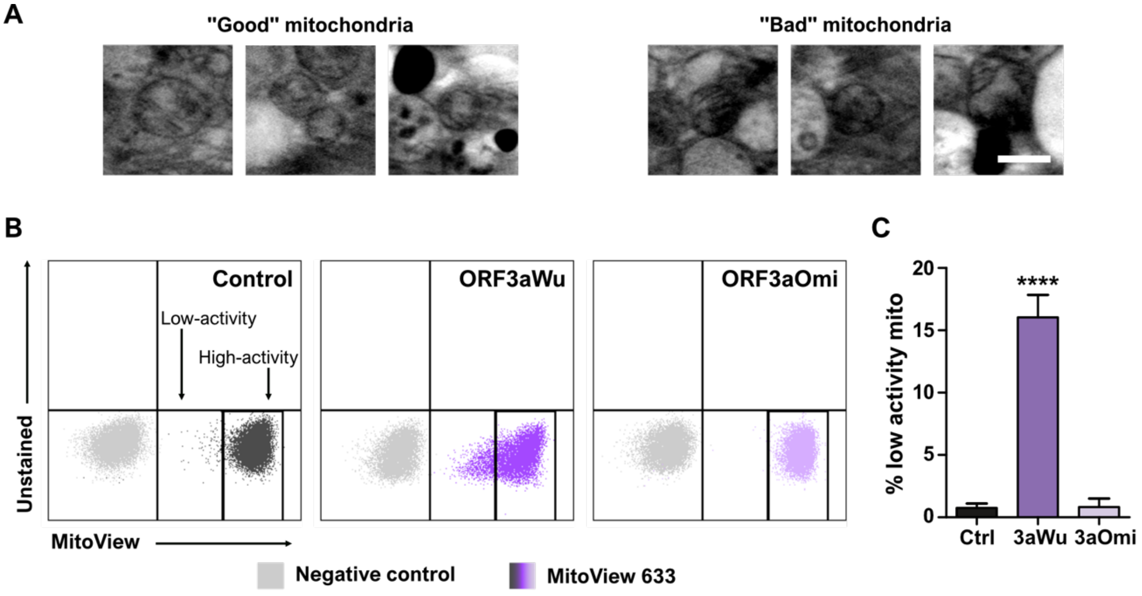


**Figure S3. Analysis of mitochondria structure and activity in A549 transduced cells. (A)** Representative cryo-SXT images of morphological unaltered ("good") and abnormal ("bad") mitochondria in ORF3aWu-expressing cells (scale bar: 1 µm). **(B)** Flow cytometry dot plots of control, ORF3aWu and ORF3aOmi-expressing cells stained with MitoView 633 (5 nM). Two cell populations were identified based on mitochondrial membrane potential: cells with high and low mitochondrial activity. **(C)** Quantification of the percentage of cells with low mitochondrial activity in control, ORF3aWu- and ORF3aOmi-expressing cells stained with MitoView 633. Data are presented as mean ± SD (n = 3). Statistical significance is as follows: ****$p < 0.0001$.

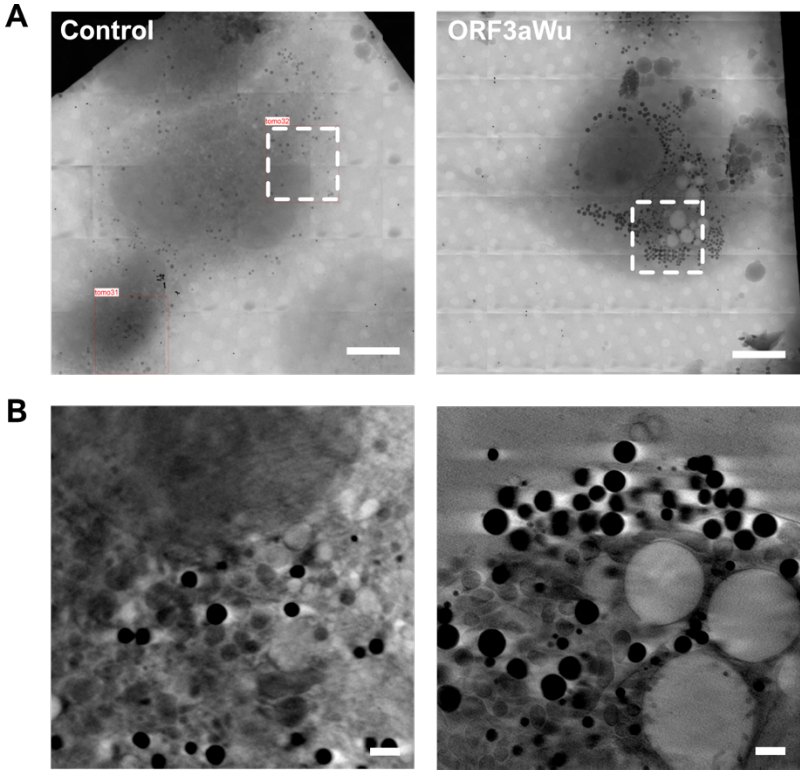


**Figure S4. THP-1 cells cryo-SXT. (A)** 3D reconstruction of control and ORF3aWu-expressing THP-1-PMA differentiated cells by cryo-SXT. Top panel full view from a single cell (scale bar: 10 μm). Dashed boxes indicate regions of tomogram acquisition (**(B)**, scale bar: 1 μm). Bottom panels show the corresponding tomograms.

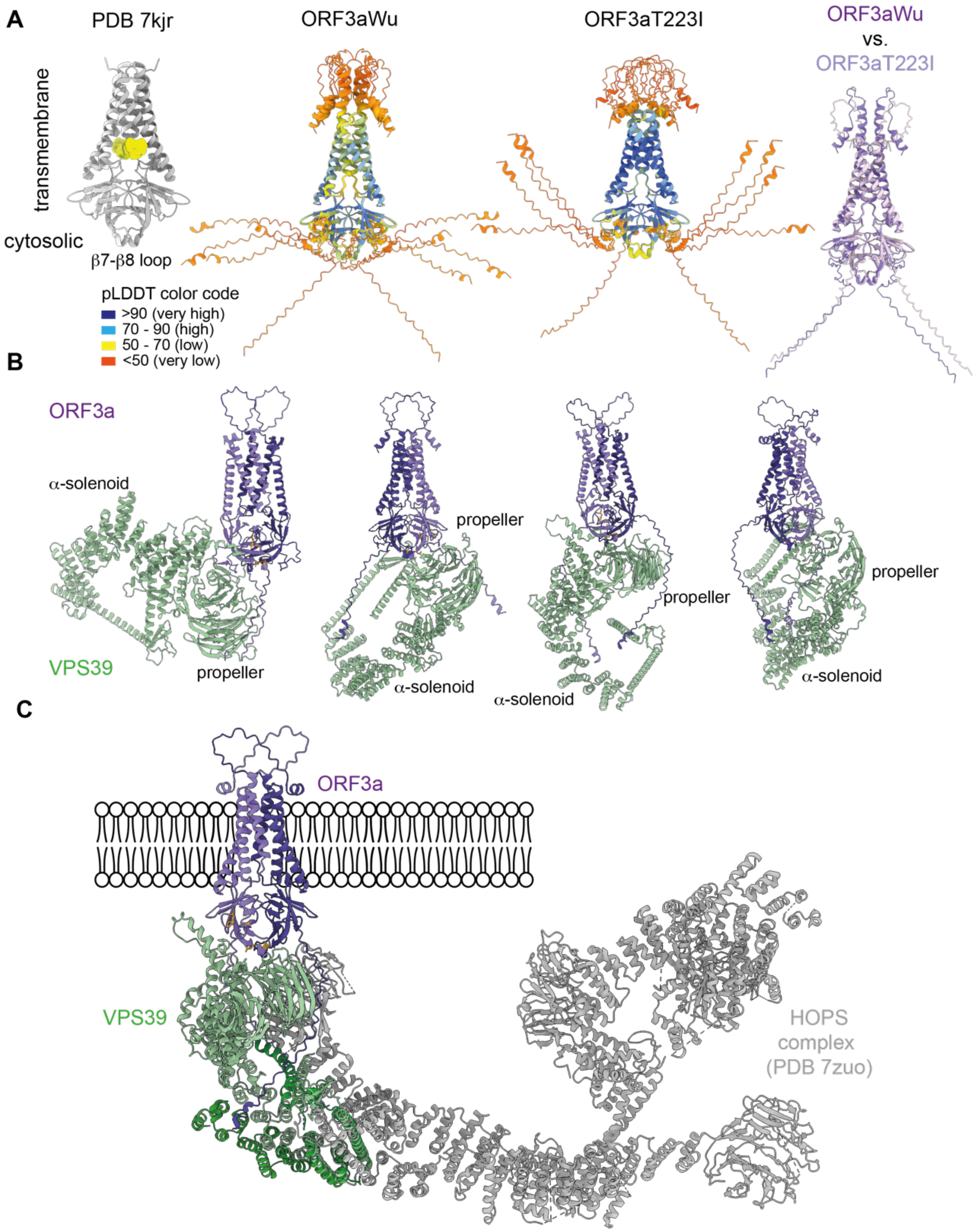


**Figure S5**: **AlphaFold3 structural models of ORF3a and its interaction with VPS39. (A)** Comparison of the experimental cryo-EM structure of SARS-CoV-2 ORF3a (PDB 7kjr) with the highest-ranked AlphaFold3 predictions of ORF3aWu and ORF3aT223I. Models are coloured according to the predicted local distance difference test (pLDDT) confidence score. Superposition of the two AlphaFold3 models highlights the pronounced reorientation of the intrinsically disordered N-terminal luminal region induced by the T223I substitution, despite the absence of detectable structural changes around residue 223 or within the β7-β8 loop. **(B)** Representative AlphaFold3 predictions of the ORF3a-VPS39 complex. In all top-ranked models, ORF3a (purple) is predicted to interact primarily with the β-propeller domain of VPS39 (green), whereas the Rab7-binding α-solenoid region remains accessible, suggesting that ORF3a does not compete directly with Rab7 for VPS39 binding. Several predicted interfaces are consistent with previous experimental evidence identifying the ORF3a β2-β3 loop as a major determinant of VPS39 interaction. **(C)** Structural model illustrating the predicted positioning of ORF3a on VPS39 within the context of the HOPS complex. The ORF3a-VPS39 complex corresponds to an AlphaFold3 prediction, whereas the HOPS complex is represented by the experimentally determined structure (PDB 7zuo). The model suggests that ORF3a associates with VPS39 without sterically occluding the Rab7-binding α-solenoid, supporting a mechanism whereby ORF3a modulates HOPS function through reorientation of VPS39 rather than direct competition for Rab7 binding.

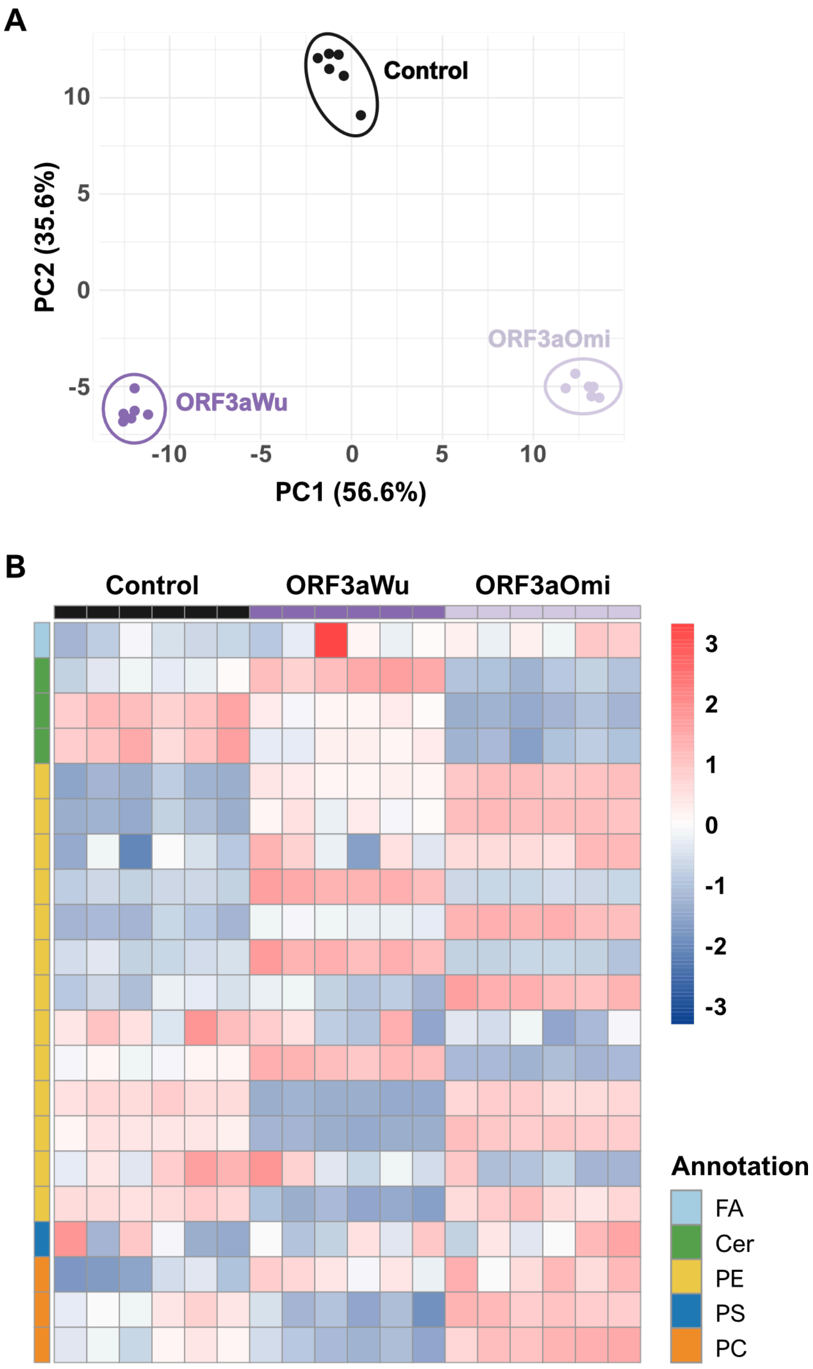


**Figure S6. Lipidomic analysis. (A)** Principal Component Analysis (PCA) graph showing distinct clustering of lipidomic profiles of control, ORF3aWu and ORF3aOmi-transduced A549 cells. **(B)** Heatmap showing altered lipids obtained in negative ionization mode in control, ORF3aWu and

ORF3aOmi A549 cells, sorted by lipid families: ceramides (Cer), phosphatidylcolines (PC), phosphatidyletanolamines (PE), fatty acyls (FA) and phosphatidylserines (PS). Color key corresponds to row-normalized values of lipid concentration.

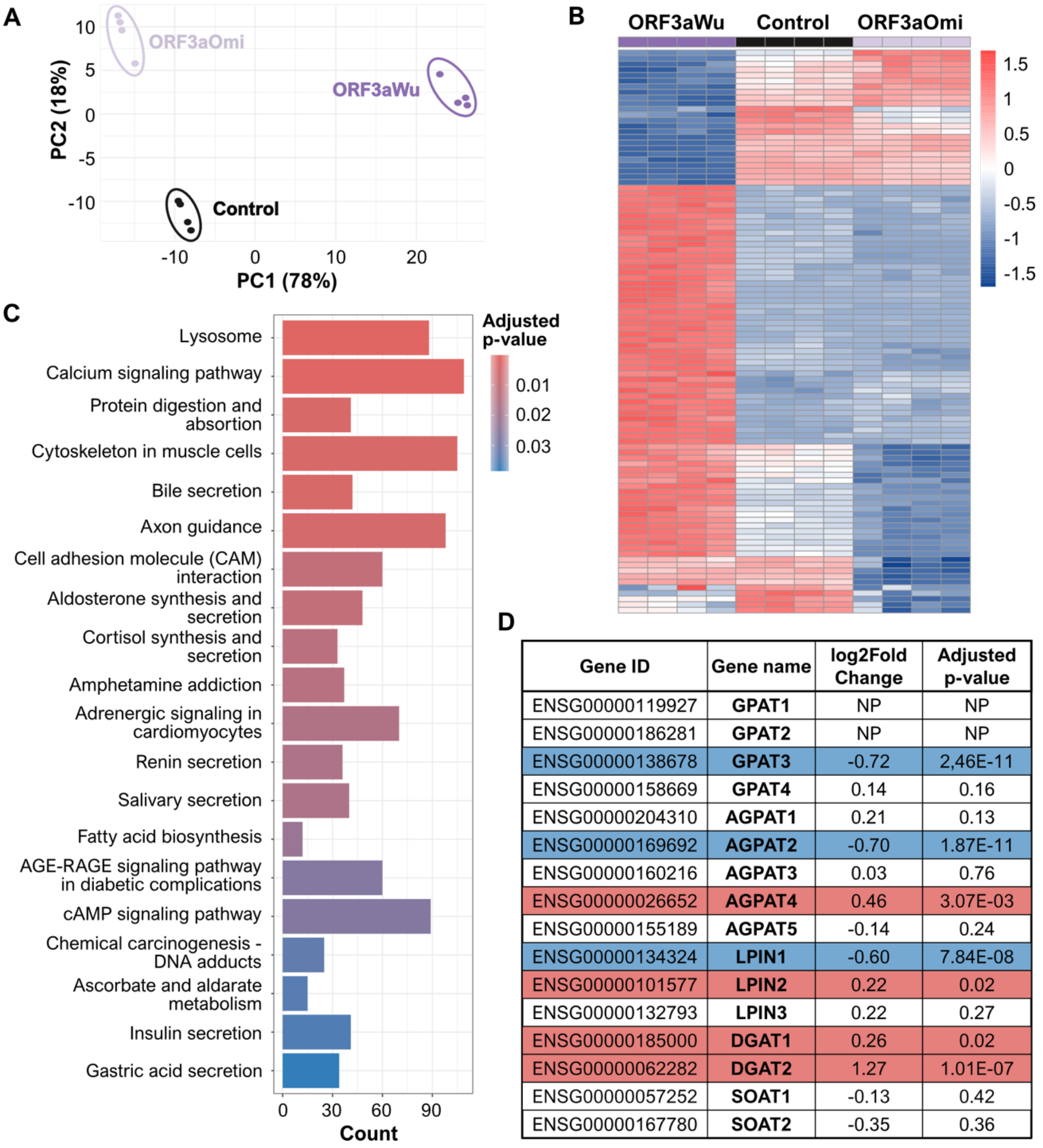


| Gene ID | Gene name | log2Fold Change | Adjusted p-value |
|---|---|---|---|
| ENSG00000119927 | **GPAT1** | NP | NP |
| ENSG00000186281 | **GPAT2** | NP | NP |
| ENSG00000138678 | **GPAT3** | -0.72 | 2,46E-11 |
| ENSG00000158669 | **GPAT4** | 0.14 | 0.16 |
| ENSG00000204310 | **AGPAT1** | 0.21 | 0.13 |
| ENSG00000169692 | **AGPAT2** | -0.70 | 1.87E-11 |
| ENSG00000160216 | **AGPAT3** | 0.03 | 0.76 |
| ENSG00000026652 | **AGPAT4** | 0.46 | 3.07E-03 |
| ENSG00000155189 | **AGPAT5** | -0.14 | 0.24 |
| ENSG00000134324 | **LPIN1** | -0.60 | 7.84E-08 |
| ENSG00000101577 | **LPIN2** | 0.22 | 0.02 |
| ENSG00000132793 | **LPIN3** | 0.22 | 0.27 |
| ENSG00000185000 | **DGAT1** | 0.26 | 0.02 |
| ENSG00000062282 | **DGAT2** | 1.27 | 1.01E-07 |
| ENSG00000057252 | **SOAT1** | -0.13 | 0.42 |
| ENSG00000167780 | **SOAT2** | -0.35 | 0.36 |

**Figure S7. Transcriptomic analysis. (A)** PCA graph of the RNA-sequencing (RNA-Seq) data from control, ORF3aWu and ORF3aOmi-transduced A549 cells. **(B)** Heatmap of RNA-Seq analysis of A549 control, ORF3aWu and ORF3aOmi cells. **(C)** Bar plot showing overrepresentation analysis of enriched KEGG pathways in ORF3aWu vs control A549 cells. Count indicates number of differentially expressed genes in each KEGG pathway. **(D)** Table showing differentially expressed genes (adjusted p-value<0.05) involved in lipid droplet biogenesis pathway in ORF3aWu vs control A549 cells. Red and blue colours indicate upregulated and downregulated genes respectively according to the adjusted. NP: not present.

**Table S1**. **List of primers used for real time qPCR.**

| Target gene | Forward primer (5'-3') | Reverse primer (5'-3') | Reference |
|---|---|---|---|
| **2xST** | TTTGAGAAGGGTGGTGGTTC | GCAGCGCCTTTTTCAAATTG C | This work |
| **ORF3aWu-2xST** | GTACGCGCTCGTTTACTTCC | GAGGATTCTTGCTCCGACAC | López-Ayllón et al., 2024 |
| **ORF3aOmi-2xST** | GTACGCGCTCGTTTACTTCC | GAGGATTCTTGCTCCGACAC | López-Ayllón et al., 2024 |
| **eGFP** | AAGGACGACGGCAACTACA A | CGATGTTGTGGCGGATCTTG | This work |
| **ORF3aWu-GFP** | GATGAGCCAGAGGAGCATG T | TGAACTTCAGGGTCAGCTTG C | This work |
| **ORF3aOmi-GFP** | GATGAGCCAGAGGAGCATG T | TGAACTTCAGGGTCAGCTTG C | This work |
| **DGAT1** | GCTTCAGCAACTACCGTGGC AT | CCTTCAGGAACAGAGAAAC CACC | Longo et al., 2024 |
| **DGAT2** | TCCAGCTGGTGAAGACACAC | GCTGACAGGGCAGATACCTC | Longo et al., 2024 |
| **GAPDH** | TGGGTGTGAACCATGAGAA G | TGGCAGTGATGGCATGGAC | López-Ayllón et al., 2024 |

**Table S2**. **List of antibodies used for immunofluorescence assays (IF) and Western Blot (WB)**.

| Antibody | Supplier | Catalog number | Assay |
|---|---|---|---|
| Chicken anti-GFP | Abcam | #ab13970 | IF |
| Goat anti-chicken-IgG (H+L) Alexa Fluor™ 488 | Invitrogen | #A11039 | IF |
| Rabbit anti-Golgi (Giantin) | Proteintech | #222701AP | IF |
| Goat anti-rabbit-IgG (H+L) highly cross-adsorbed Alexa Fluor™ 546 | Invitrogen | #A11035 | IF |
| Mouse anti-human Cd63 | Kindly provided by Dr. F Sanchez-Madrid | clone TEA3/18 | IF |
| Goat anti-Mouse-IgG (H+L) highly cross-adsorbed Alexa Fluor 647™ | Invitrogen | #A21236 | IF |
| Rabbit anti-DGAT1 | Selleckchem | #F3308 | WB |
| Rabbit anti-DGAT2 | ThermoFisher Scientific | #PA5103785 | WB |
| Mouse anti-GAPDH | Sigma | #G8795 | WB |
| Goat anti-rabbit-IgG-HRP | Abcam | #ab6721 | WB |
| Rabbit anti-mouse-IgG-HRP | Sigma | #A9044 | WB |
